# Genotype & Phenotype in Lowe Syndrome: Specific *OCRL1* patient mutations differentially impact cellular phenotypes

**DOI:** 10.1101/2020.08.04.236612

**Authors:** Swetha Ramadesikan, Lisette Skiba, Jennifer Lee, Kayalvizhi Madhivanan, Daipayan Sarkar, Agustina De La Fuente, Claudia B. Hanna, Genki Terashi, Tony Hazbun, Daisuke Kihara, R. Claudio Aguilar

## Abstract

Lowe Syndrome (LS) is a lethal genetic disorder caused by mutations in the *OCRL1* gene which encodes the lipid 5’ phosphatase Ocrl1. Patients exhibit a characteristic triad of symptoms including eyes, brain and kidneys abnormalities with renal failure as the most common cause of premature death. Over 200 *OCRL1* mutations have been identified in LS, but their specific impact on cellular processes is unknown. Despite observations of heterogeneity in patient symptom severity, there is little understanding of the correlation between genotype and its impact on phenotype.

Here, we show that different mutations had diverse effects on protein localization and on triggering LS cellular phenotypes. In addition, some mutations affecting specific domains imparted unique characteristics to the resulting mutated protein. We also propose that certain mutations conformationally affect the 5’-phosphatase domain of the protein, resulting in loss of enzymatic activity and causing common and specific phenotypes.

This study is the first to show the differential effect of patient 5’-phosphatase mutations on cellular phenotypes and introduces a conformational disease component in LS. This work provides a framework that can help stratify patients as well as to produce a more accurate prognosis depending on the nature and location of the mutation within the *OCRL1* gene.

## INTRODUCTION

Lowe Syndrome (LS) or Oculo-Cerebro-Renal syndrome of Lowe (OCRL) (OMIM#30900) is an X-linked genetic disorder caused by mutations in the *OCRL1* gene (1). Affected children are born with bilateral cataracts, present neurological abnormalities and renal symptoms a few months after birth (2, 3). Progressive renal dysfunction leads to end stage renal disease and premature death (2). Unfortunately, currently there are no LS-specific therapeutics (4).

The gene product of *OCRL1* is the inositol 5’ phosphatase Ocrl1 (EC 3.1.3.36) which has specificity for the signaling lipid phosphatidyl inositol 4,5-bisphophate, PI(4,5)P2 (5, 6). Ocrl1 localizes at the *trans*-Golgi network (TGN) (7), endosomes (8) and transiently at the plasma membrane (9). In addition to its phosphatase domain, Ocrl1 possesses N-terminal PH (10) and C-terminal ASH-RhoGAP (8, 11) domains through which it interacts with several signaling and trafficking proteins; thereby participating in several basic cellular processes including cell migration, cell spreading, actin remodeling, ciliogenesis, vesicle trafficking, cytokinesis and phagocytosis (12–22). Further, we previously established that Ocrl1 participates in certain processes through determinants spatially segregated within the protein. Specifically, that the N-terminus is required for membrane remodeling (12) and the C-terminus for primary cilia assembly (15)); while a functional Ocrl1 5’-phosphatase domain is required for both processes to proceed normally (12, 15).

There are over 200 unique LS-causing mutations that have been identified in *OCRL1* (4, 23) affecting different domains of Ocrl1. Therefore, it could be expected that mutations affecting specific regions of Ocrl1, would have a differential impact on cellular phenotypes causing heterogeneity in LS patient manifestations; however, little is known about the correlation between the genotype of patients and their cellular phenotypes.

Here we show that depending on the domain they affect, *OCRL1* mutations have a differential impact on cell spreading and ciliogenesis, thereby suggesting cellular basis for LS patient symptom heterogeneity. In addition, specific mutants possess unique characteristics in terms of localization and protein stability. Importantly, some patient variants bearing a mutated phosphatase domain induced fragmentation of the Golgi apparatus. This phenotype was not induced by any other patient mutation tried. This cellular phenotype, although novel for LS, it has been previously observed in diseases affecting the nervous system (24–26). Therefore, and given that LS has neurological manifestations, we speculate that this defect may play a role in disease pathogenesis.

In addition, molecular dynamics analysis of Ocrl1 patient variants *with residue changes at non-catalytic sites* within the phosphatase domain predicted the existence of conformational changes affecting the active site. Further, experimental results showed impairment of 5’-phosphatase activity for these mutants as well as the presence of cellular phenotypes in cells expressing these variants. These results explain how mutations affecting non-catalytic residues cause LS, but also imply that some *OCRL1* mutants lead to a conformational/protein misfolding disease scenario.

We believe that this study will provide the framework that will allow LS patient stratification as well as aid in creating tailored therapeutic strategies that would consider the nature and location of mutation in the gene as well as its effect on the biochemical activity of Ocrl1.

## RESULTS

### Different *OCRL1* missense patient mutations affect various Ocrl1 domains

This study is focused on understanding the consequences of *OCRL1* missense mutations on several characteristics (stability, localization and ability to sustain specific cellular functions) of the encoded 5’-phosphatase Ocrl1.

Since 90% of missense mutations found in LS patients locate within *OCRL1’s* exons encoding for the phosphatase and ASH (ASPM-SPD-2-Hydin)-RhoGAP (RhoGTPase Activating Protein) domains (4, 23, 27–30), we selected patient missense mutations located within exons 9-23 (Table I and Fig. 1A).

**Table I:** *OCRL1* patient mutants used in this study

| Table I: *OCRL1* patient mutants used in this study |  |  |
| --- | --- | --- |
| Mutation | Exon | Domain affected |
| S256N | 9 | 5' phosphatase |
| D451G | 13 |  |
| V508D | 15 |  |
| H524R |  |  |
| V577E | 17 | ASH-RhoGAP |
| F668V | 18 |  |
| I768N | 21 |  |

**Fig. 1.**
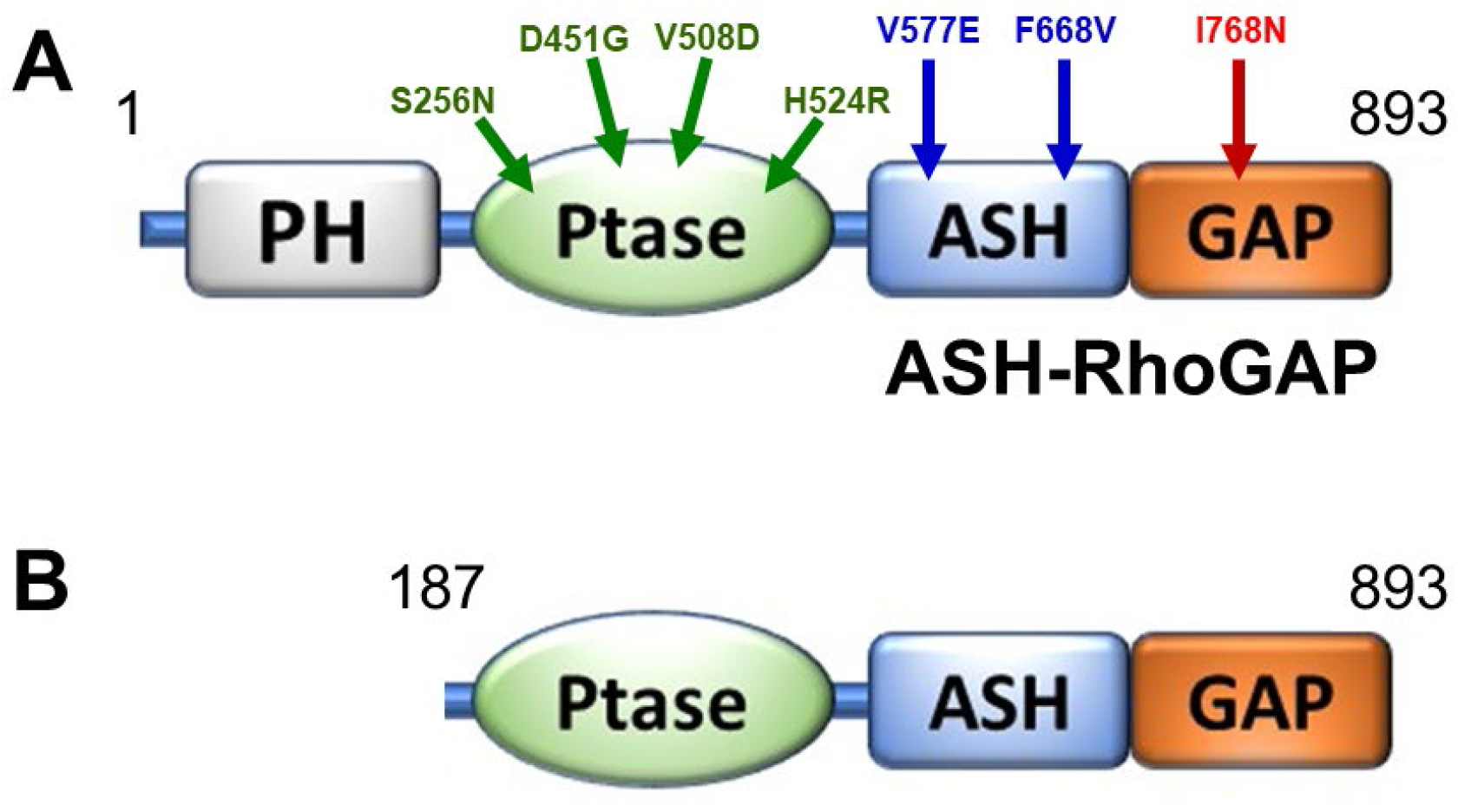
Ocrl1 variants used in this study. **A.** Residue change due to LS patient missense mutations are mapped on the Ocrl1 molecule. **B**. Representation of a proposed Ocrl1 variant resulting from alternative initiation at Met^187^. PH: Pleckstrin homology; Ptase: Inositol phosphatase domain; ASH: ASPM-SPD-2-Hydin; RhoGAP: RhoGTPase Activating Protein.

There are nearly 80 unique missense, LS-causing mutations which lead to amino acid changes affecting the Ocrl1’s phosphatase domain. While many of these replacements map to conserved regions important for catalysis and substrate recognition, others affected residues not directly involved in substrate binding and processing (Supplemental Fig. 1). Therefore, we selected Ocrl1 patient variants to represent both categories; *i.e*., H524R (critical catalytic residue replaced), S256N, D451G and V508D (affecting non-catalytic residues) (Supplemental Fig. 1). In addition, there are nearly 20 unique missense mutations resulting in changes in the amino acids encoding the C-terminal ASH-RhoGAP domain of Ocrl1. From these, we selected variants V577E, F668V and I768N (Table I and Fig. 1A).

Although a few Ocrl1 variants have been found to display amino acid changes in the PH domain, there are no reported LS-causing missense mutations in exons 2-7 that we know of. Nevertheless, several nonsense, frameshift mutations and deletions have been found within these 5’ exons of the *OCRL1* gene. Strikingly, these mutations often cause a milder condition known as Dent-2 disease (27). Interestingly, Shrimpton and co-investigators (31) proposed that in these cases an alternative initiation codon (corresponding to Ocrl1’s Met^187^) is used to translate an N-terminal truncation (lacking the PH domain) that would retain substantial functionality. We recreated such Ocrl1^187-901^ (ΔPH) product for characterization in this study (Fig.1B).

### Ocrl1 patient variants differentially affect cell spreading and ciliogenesis in human kidney epithelial cells

Since renal failure is the main cause of death among LS patients, in this study we mostly used human embryonic kidney 293T epithelial cells lacking *OCRL1* (HEK293T KO). We also focused on cell spreading and ciliogenesis as cellular readouts as they are likely to impact organogenesis and renal function.

#### Cell spreading

We transfected the different GFP-tagged Ocrl1 mutated variants or Ocrl1^WT^ in HEK293T KO cells and performed standard spreading assays as described before (12) and in *Materials and Methods*. Briefly, 18h after transfection, cells were lifted and allowed to attach and spread for 30min on fibronectin-coated coverslips. After 30min of attachment and spreading, cells were fixed with 4% formaldehyde, stained with rhodamine-phalloidin to label the actin cytoskeleton. Random fields containing transfected cells were imaged and the spreading area was measured using the *magic wand* tracing tool in ImageJ, fraction of cells vs cell area histograms were constructed and statistical analysis was performed as described in *Materials and Methods*.

In agreement with the hypothesis that Ocrl1’s N-terminus region is required for membrane remodeling (12), Ocrl1^ΔPH^-expressing cells showed a significant spreading defect detected as the corresponding histogram shifts towards smaller areas as compared to the Ocrl1^WT^ histogram (Fig 2A).

**Fig. 2.**
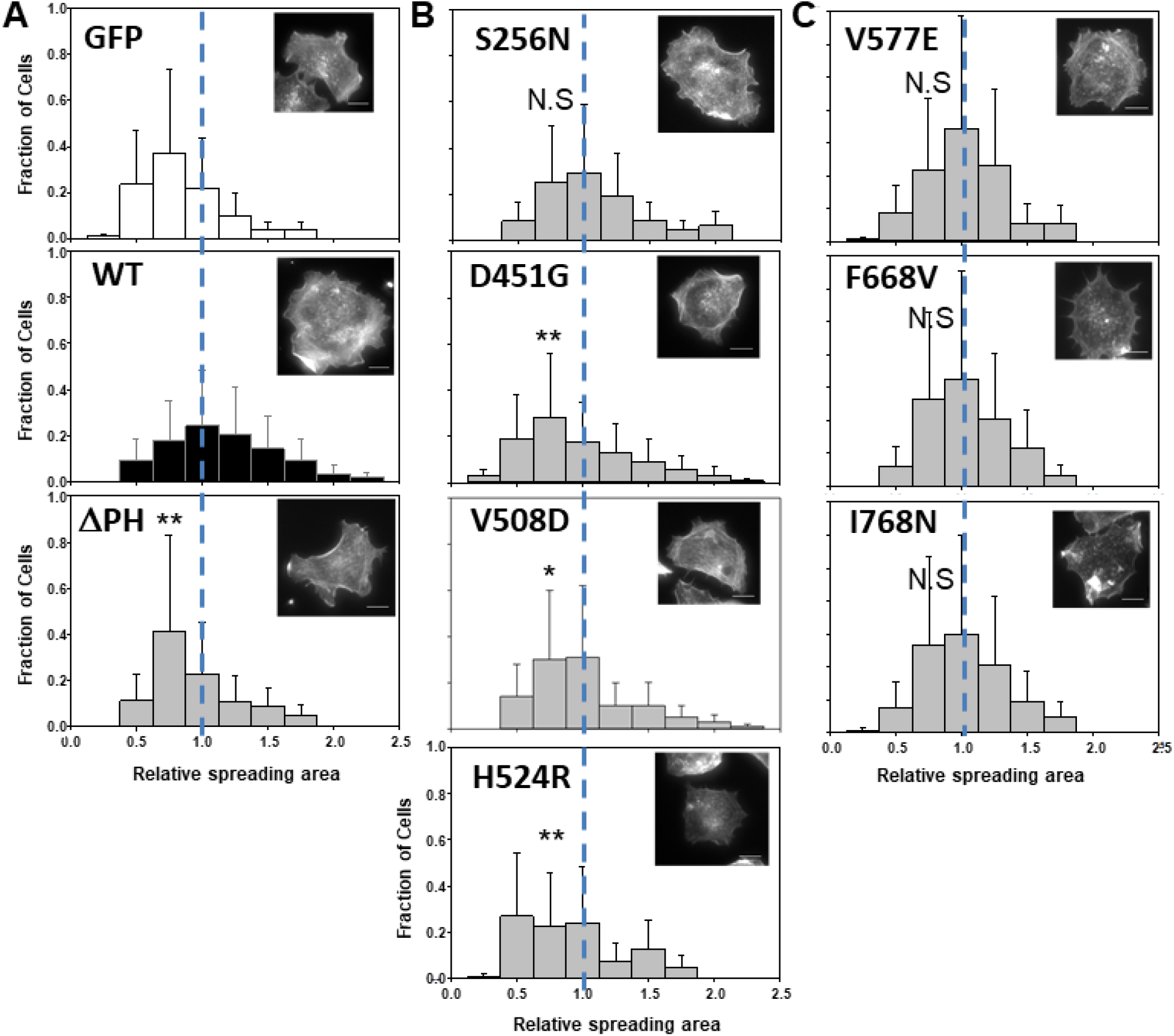
Ocrl1 patient variants differentially affect cell spreading. HEK293T KO cells were transfected with Ocrl1^WT^, Ocrl1^ΔPH^ (**A**); phosphatase domain mutants (**B**); or ASH-RhoGAP domain mutants (**C**) and allowed to attach and spread on fibronectin-coated surfaces (See *Materials and Methods*). Median spreading areas of cells expressing mutants were normalized with respect to Ocrl1^WT^. Histograms are from three independent experiments. Experiments were repeated at least 3 times, with a total n=120-150 cells. Example of rhodamine-phalloidin stained cells representative of the high frequency groups within each histogram. Scale bar: 10μm. Statistically significance of the mean difference with respect to Ocrl1^WT^ was A: **p<0.05 by KS test B: **p<(0.05/4=0.0125), *p<(0.1/4=0.025) (Bonferroni correction) by KS test, N.S not significant C: N.S not significant.

We found that cells expressing most Ocrl1 patient variants bearing a mutated phosphatase domain also displayed significant smaller spreading areas compared to those expressing Ocrl1^WT^ (Fig.2B). However, Ocrl1^S256N^ did not induce a significant spreading defect in cells (Fig.2B). Mutated patient variants within this category induced different degree of severity in the cell spreading phenotype; *e.g*., while Ocrl1^D451G^ and Ocrl1^H524R^ showed a more severe spreading phenotype, Ocrl1^V508D^ was only modestly affected (~12% reduction in median spreading area compared to WT) and as indicated, Ocrl1^S256N^ produced no spreading phenotype (Fig.2B).

Although cells expressing ASH-RhoGAP mutant Ocrl1^V577E^, Ocrl1^F668V^, Ocrl1^I768N^ were somewhat prone to show smaller cell spreading area, the differences were not statistically significant as compared to WT (Fig.2C). These results were in agreement with previous observations that indicated that the C-terminal region of Ocrl1 was not critical for membrane remodeling (12).

#### Ciliogenesis

We also tested the impact of Ocrl1 patient mutated variants on primary cilia (PC) assembly, to such effect HEK293T KO cells were seeded at ~30-50% confluency, transfected with plasmids for expression of Ocrl1 WT/patient variants and ciliogenesis assays were performed as described previously (15) and in *Materials and Methods*. Briefly, 18h after transfection, cells were starved for 24h by maintaining them in 0.1% FBS media to induce ciliogenesis. After that, cells were fixed with 4% formaldehyde, followed by indirect immunofluorescence using antibodies against acetylated tubulin (to label primary cilia) and pericentrin-2 (to label centriole at the base of cilia). Fields were randomly imaged and the fraction of Ocrl1 mutant transfected cells forming cilia was determined and compared to the fraction of ciliated Ocrl1^WT^-expressing cells (See *Materials and Methods*).

Interestingly, Ocrl1^ΔPH^ was unaffected for ciliogenesis (Fig.3A), likely due to the presence of functional phosphatase and ASH-RhoGAP domains which were previously shown by our lab to be required for PC assembly (15). These results also suggested that the N-terminal truncated Ocrl1 variant starting at M^187^ is partially functional; *i.e*., despite lacking the ability to facilitate cell spreading, it was capable of sustaining ciliogenesis.

**Fig. 3.**
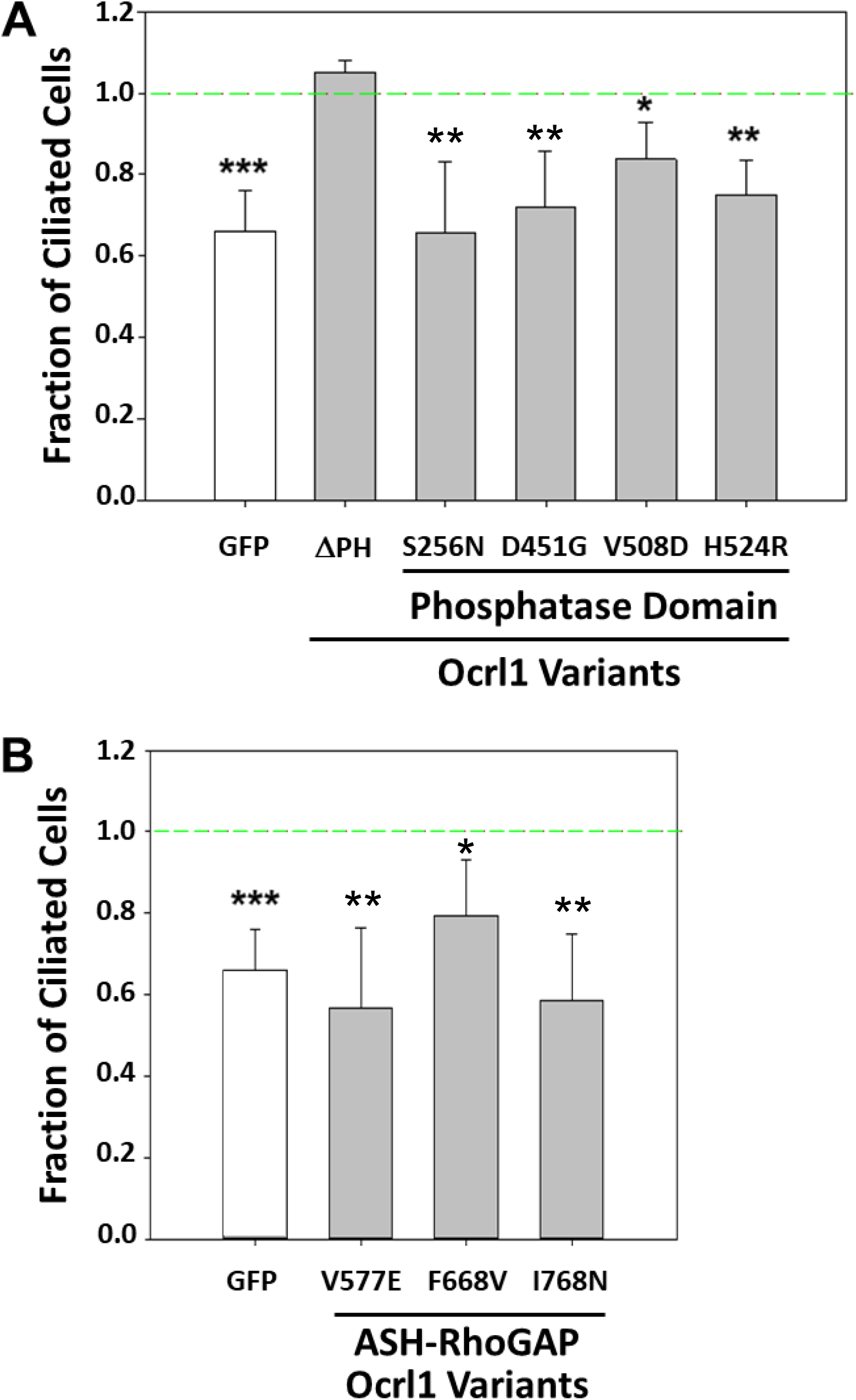
Ocrl1 patient variants differentially affect ciliogenesis. HEK293T KO cells were transfected with different Ocrl1^WT^ or Ocrl1^ΔPH^, phosphatase domain mutants (**A**) or ASH-RhoGAP domain mutants (**B**) and ciliogenesis assays were performed (see *Materials and Methods*). 20 random fields with at least 50 cells were imaged and fraction of transfected cells with cilia were calculated. This number was normalized to the fraction of Ocrl1^WT^-expressing cells forming cilia. Each experiment was repeated at least thrice (n=120-150 cells). Statistically significance of the mean difference with respect to Ocrl1^WT^ was **A**: *** p<0.01/6=0.0016, **p<(0.05/6=0.008), *p<(0.1/6=0.016) (Bonferroni correction) by Student’s t-test; **B**: **p<(0.05/4=0.0125), *p<(0.1/4=0.025) (Bonferroni correction) by the Student’s t-test. Reference line represents fraction of Ocrl1^WT^ expressing cells making cilia.

All cells expressing phosphatase mutants were affected for ciliogenesis in varying degrees (Fig.3A). Similar to their effect on the cell spreading, ciliogenesis phenotype severity varied among different patient variants. While cells expressing Ocrl1^H524R^, Ocrl1^S256N^, Ocrl1^D451G^ were clearly impaired for PC assembly, expression of Ocrl1^V508D^ caused a significant but milder phenotype.

In agreement with previous findings, indicating that Ocrl1’s C-terminus was important for cilia assembly, cells expressing the ASH-RhoGAP V577E, F668V and I768N mutated variants were affected for ciliogenesis in different degree as compared to Ocrl1^WT^ (Fig.3B).

### Cellular phenotypes and protein localization abnormalities associated with specific Ocrl1 patient mutated variants

In addition to well-established LS cellular phenotypes such as defects in membrane remodeling (*e.g*., cell spreading) and ciliogenesis, certain patient mutated variants exhibited the following specific phenotypes and/or localization abnormalities: a) *Ocrl1^ΔPH^: deficient localization*; b) *Ocrl1 phosphatase mutants: induction of Golgi apparatus fragmentation and disperse punctate pattern with poor TGN co-localization*. c) *Ocrl1 variants bearing mutated ASH-RhoGAP domain: mislocalization to centriolar structures* (some of these variants also exhibited cytosolic aggregates).

#### a) deficient localization of Ocrl1^ΔPH^

GFP-Ocrl1^ΔPH^ localized to the TGN similar to Ocrl1^WT^(Fig. 4, 2^nd^ and 3^rd^ image row), but surprisingly, it showed an increased cytosolic and nuclear localization (Fig.4, bottom panels). Since it is well-known that GFP has non-specific affinity for the nucleus (Fig.4, top left panel), we speculate that lack of the PH domain weakens Ocrl1 ability to properly localize, causing the protein to remain in the cytosol and susceptible to be dragged into the nucleus by fusion to GFP. Therefore, this aberrant mis-localization of the Ocrl1^ΔPH^ variant to the nucleus suggests that the N-terminal somehow contributes to maintain proper Ocrl1’s intracellular localization.

**Fig. 4.**
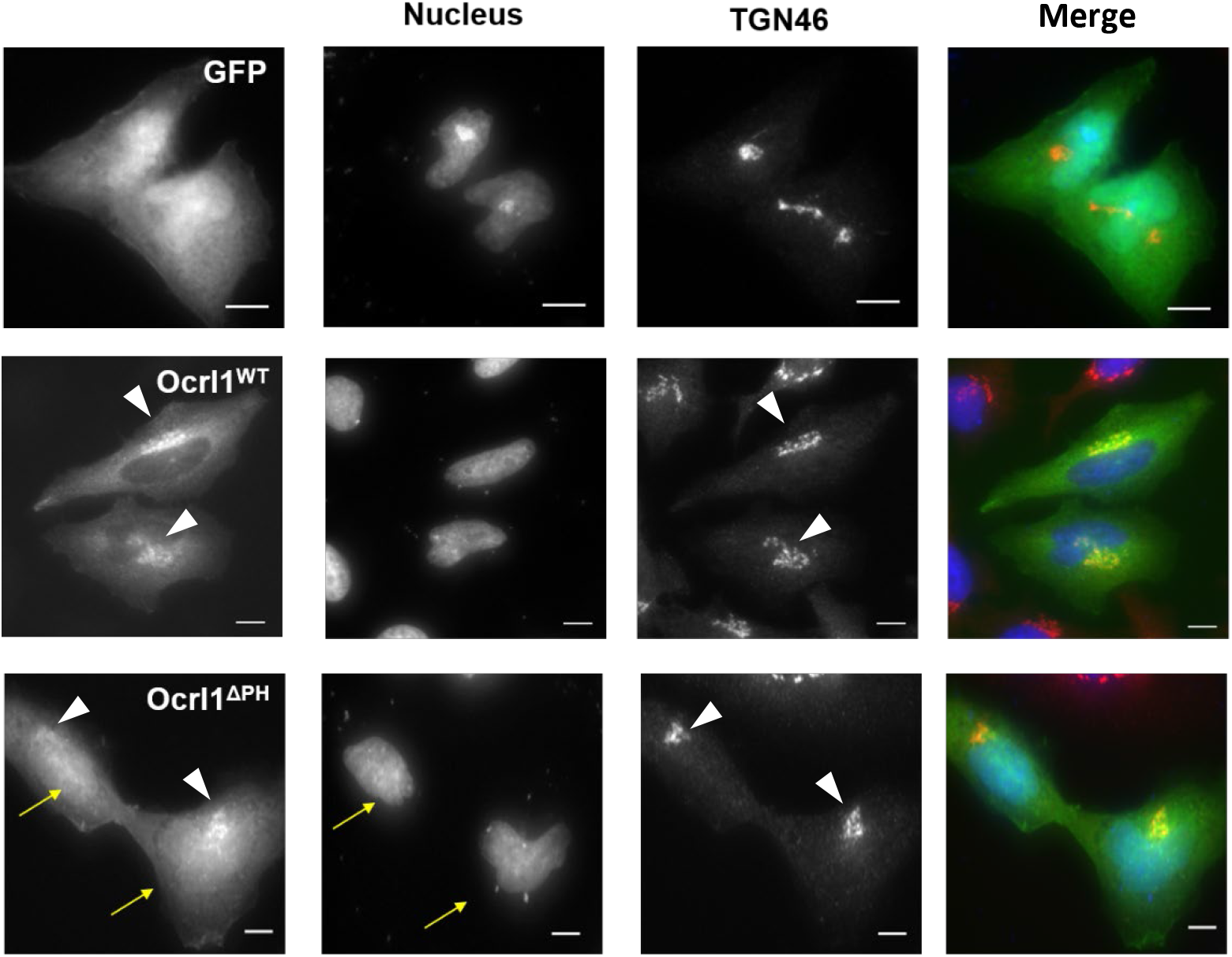
Truncation of Ocrl1 PH domain results in nuclear mislocalization of variant. HK2 KO cells transiently expressing GFP, Ocrl1^WT^ or Ocrl1^ΔPH^ and immunostained for TGN (see *Materials and Methods*). Arrows indicate Ocrl1^ΔPH^ enrichment in the nuclear compartment; arrowheads point to TGN and TGN-colocalizing Ocrl1. Scale bar: 10μm.

#### b) Induction of Golgi apparatus (GA) fragmentation

We had previously observed that overexpression of the patient’s phosphatase-dead Ocrl1^H524R^ variant led to fragmentation of the Golgi complex in HeLa cells (12). It should be highlighted that GA fragmentation has been associated with neurological disorders including Alzheimer’s, Parkinson’s and Huntington’s diseases, as well as amyotrophic lateral sclerosis, Angelman syndrome, spinal muscular atrophy and epilepsy (24–26, 32, 33). Given the presence of a neurological component in LS (2, 4), we speculated that this cellular phenotype is relevant to this disease pathogenesis. In addition, GA-related secretion defects have also been observed in polycystic kidney disease (34–36).

Since previous results (12) suggested that lack of phosphatase activity is important for this phenotype to manifest, we tested the ability of Ocrl1 patient variants bearing mutated 5’-phosphatase domains to induce GA fragmentation (measured as the ratio between the area occupied by the GA and the whole cell area, see *Materials and Methods*—Fig. 5A,B) in human kidney HK2 *OCRL1* K.O. (HK2 KO) cells. HK2 cells are larger in size and possess a flatter morphology than HEK293T, making them more suitable to microscopy analysis (in addition, all phenotypes described before for each Ocrl1 mutated variant were also observed in in HK2 KO cells (*e.g*., Figs 5, 6 and data not shown).

**Fig. 5.**
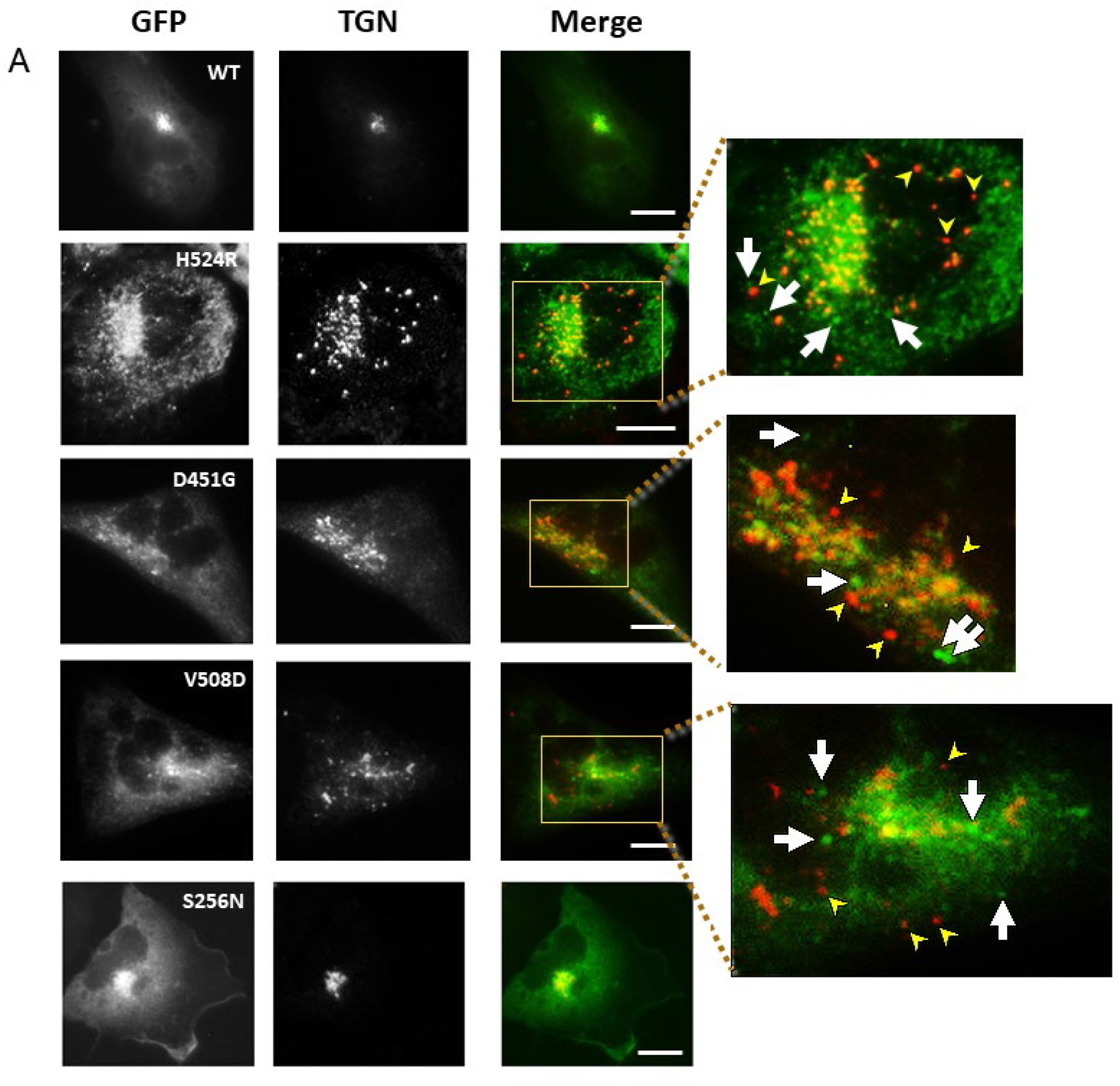
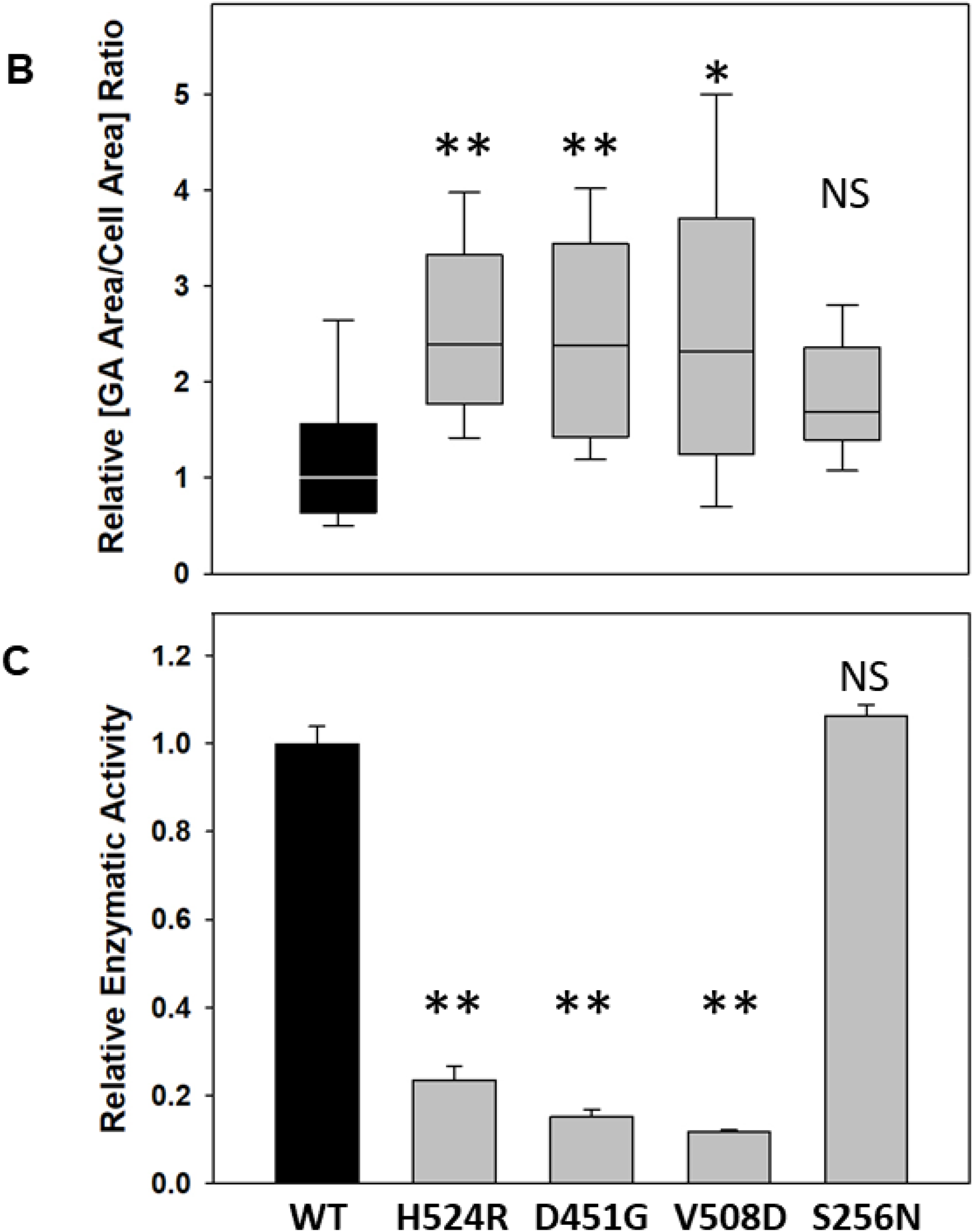
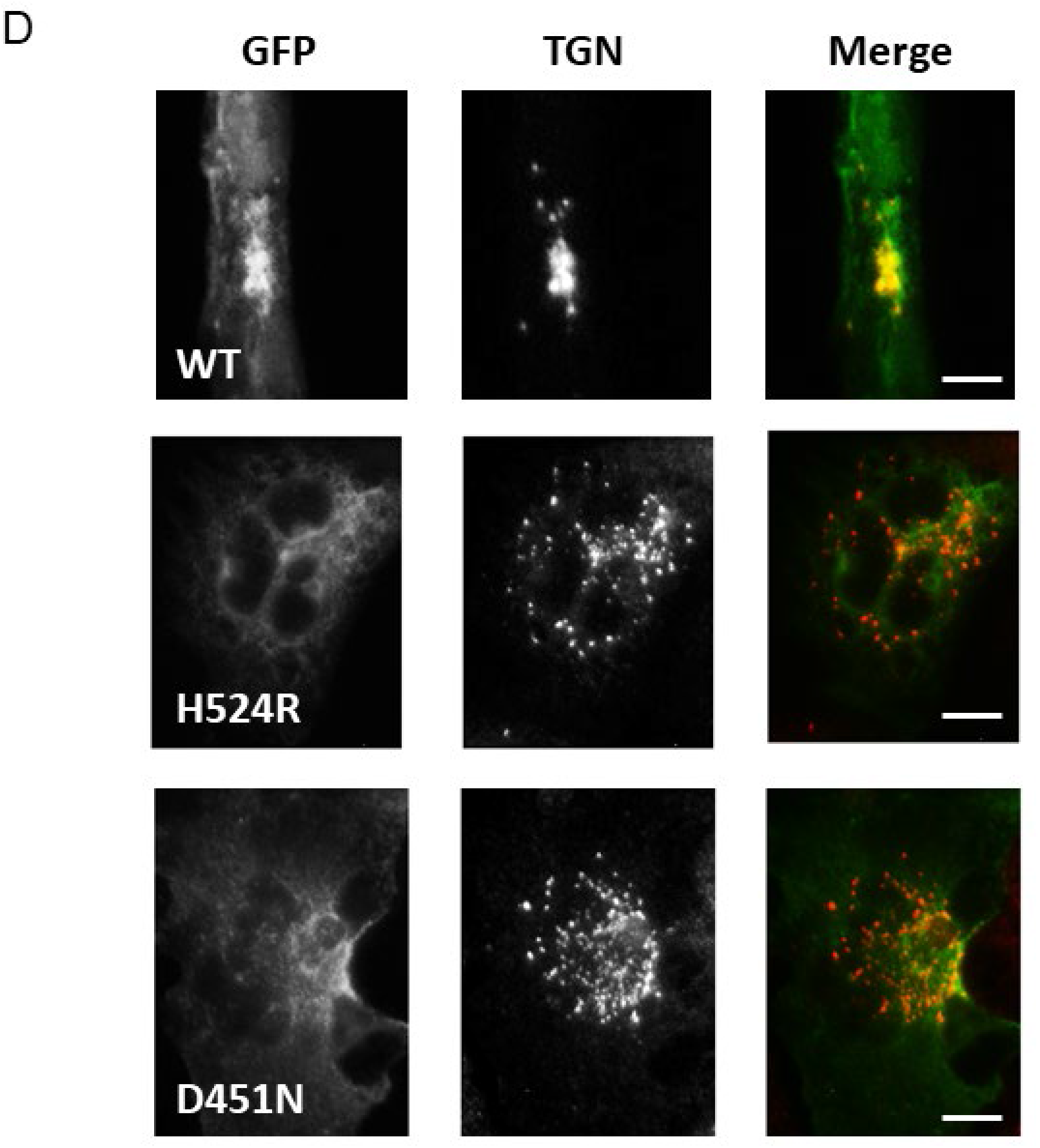
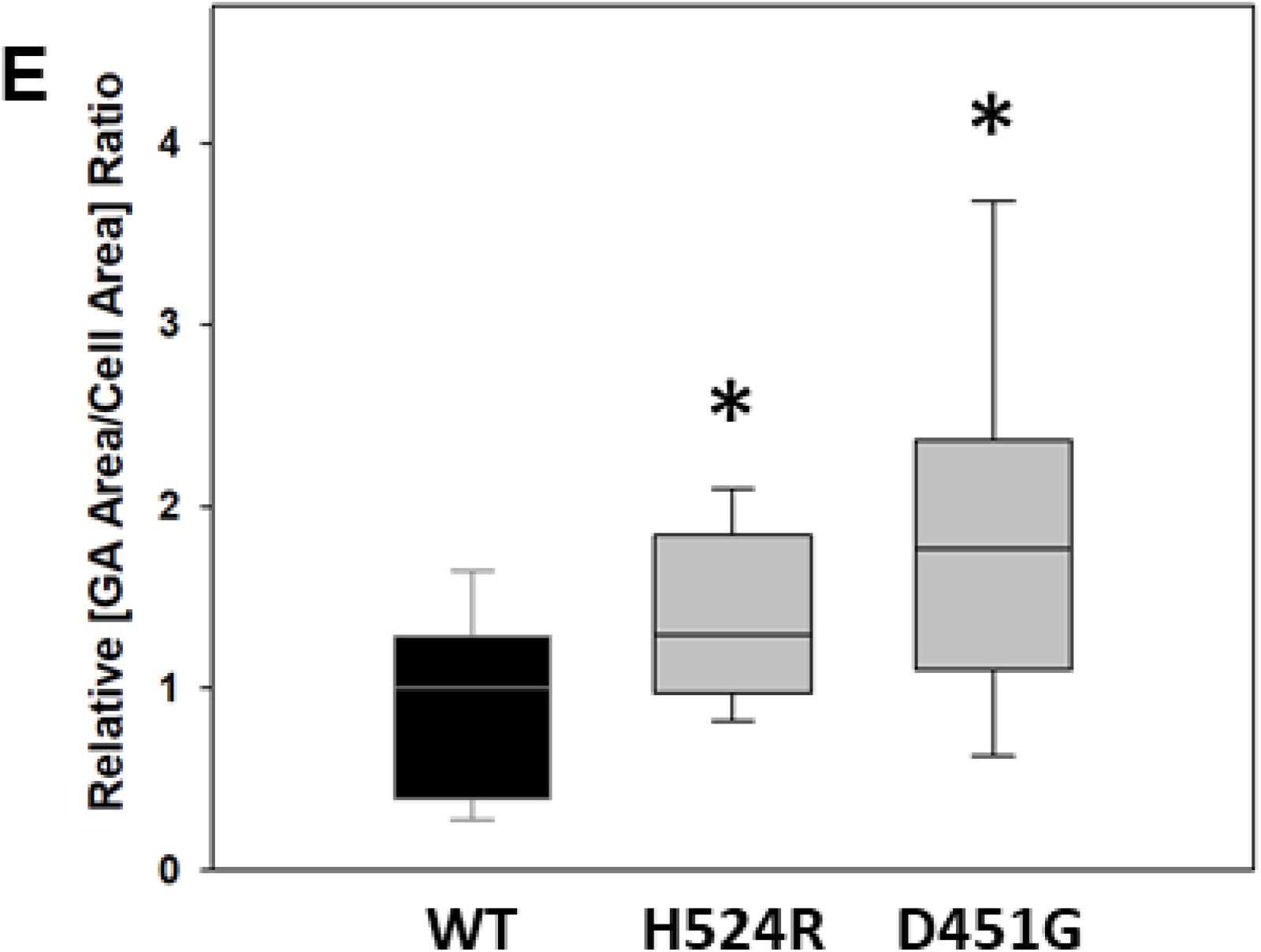
Phosphatase domain mutants produce Golgi apparatus fragmentation. **A**: HK2 KO cells transiently transfected with Ocrl1^WT^, Ocrl1^H524R^, Ocrl1^D451G^, Ocrl1^V508D^ or Ocrl1^S256N^ were immunostained for TGN. Highlighted region in merged images corresponding to the TGN area which was scaled to 3X (inset images) to better visualize Golgi apparatus fragmentation. Arrows indicate fragmented TGN puncta that lack Ocrl1. Arrowheads indicate dispersed mutant Ocrl1 lacking colocalization. Scale bar: 10μm. **B**: At least 40 transfected cells were imaged randomly, per experiment and area occupied by TGN was quantified (See *Materials and Methods*). Each experiment was repeated at least thrice with total n=120-150 cells/group. Statistically significance of the mean difference with respect to Ocrl1^WT^ was **p<(0.05/4=0.0125), *p<(0.1/4=0.025) (Bonferroni correction) by the Wilcoxon test. **C**: Golgi apparatus fragmentation plotted as a function of total fluorescence intensity of Ocrl1 (WT/patient variant) transfected in HK2 KO cells. Statistically significance of the mean difference with respect to Ocrl1^WT^ was **p<(0.05/4=0.0125) (Bonferroni correction) by the t-test. **D**: HK2 KO cells stably expressing Ocrl1^WT^ or Ocrl1^H524R^ or Ocrl1^D451G^ (see *Materials and Methods*) were prepared. All cells stably expressing Ocrl1 variants were imaged and TGN area was quantified (see *Materials and Methods*). Scale bar: 10μm. **E**: Golgi apparatus fragmentation determined as a function of total cell area. Statistically significance of the mean difference with respect to Ocrl1^WT^ was **p<(0.05/2=0.025) (Bonferroni correction) by the Wilcoxon-test.

**Fig. 6.**
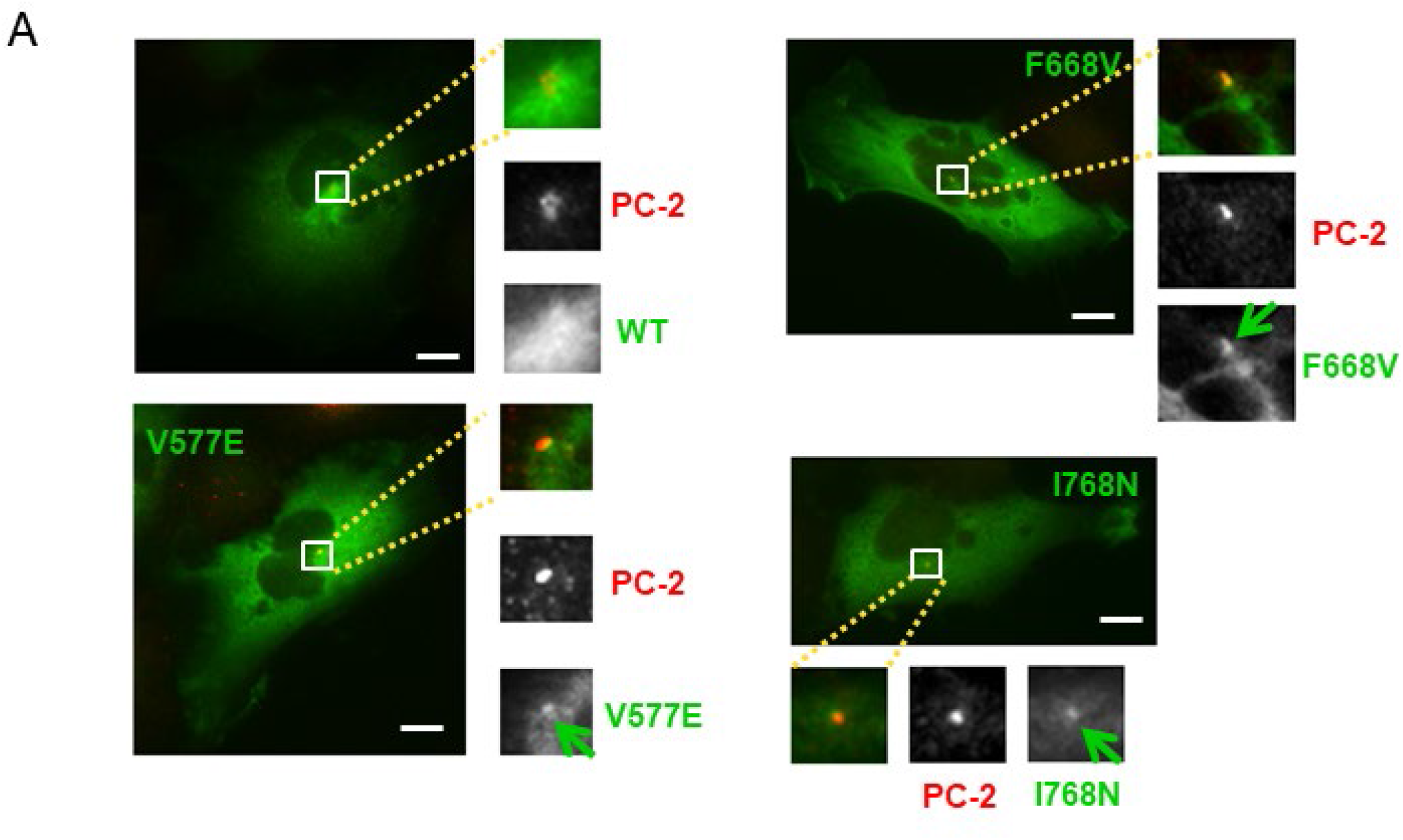
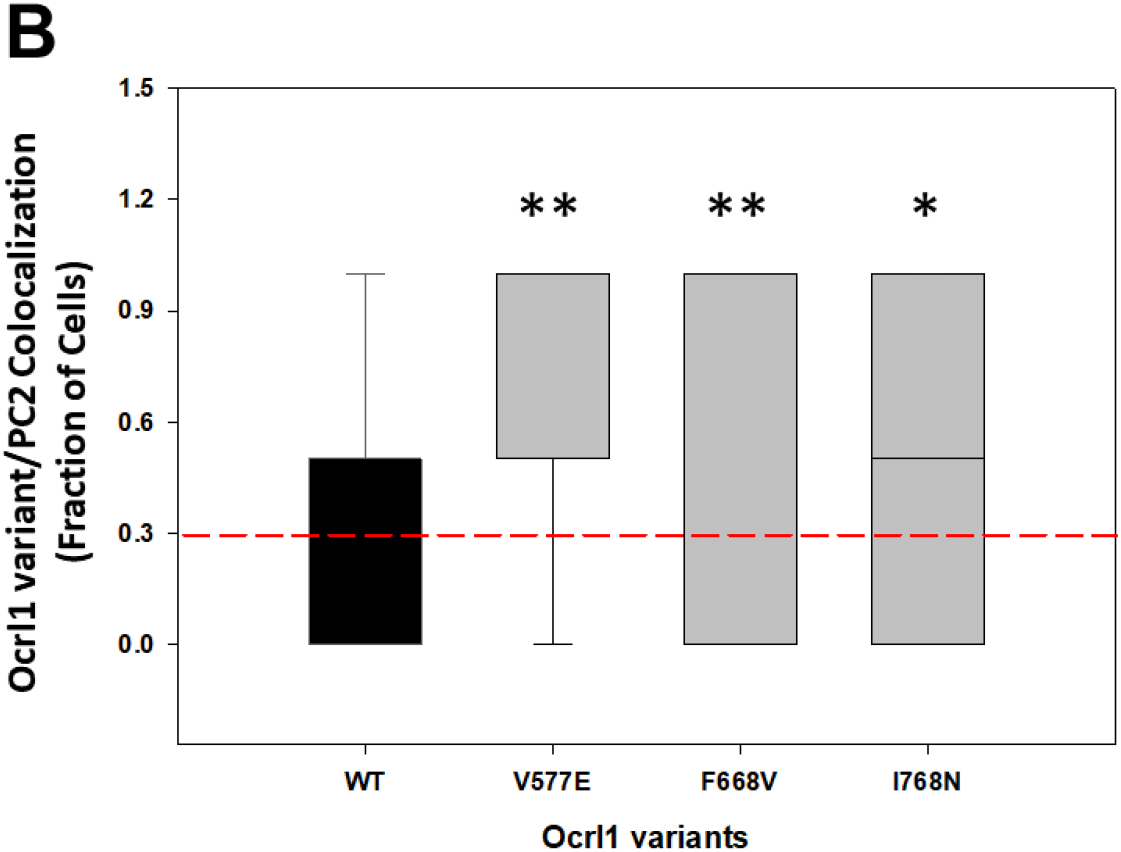
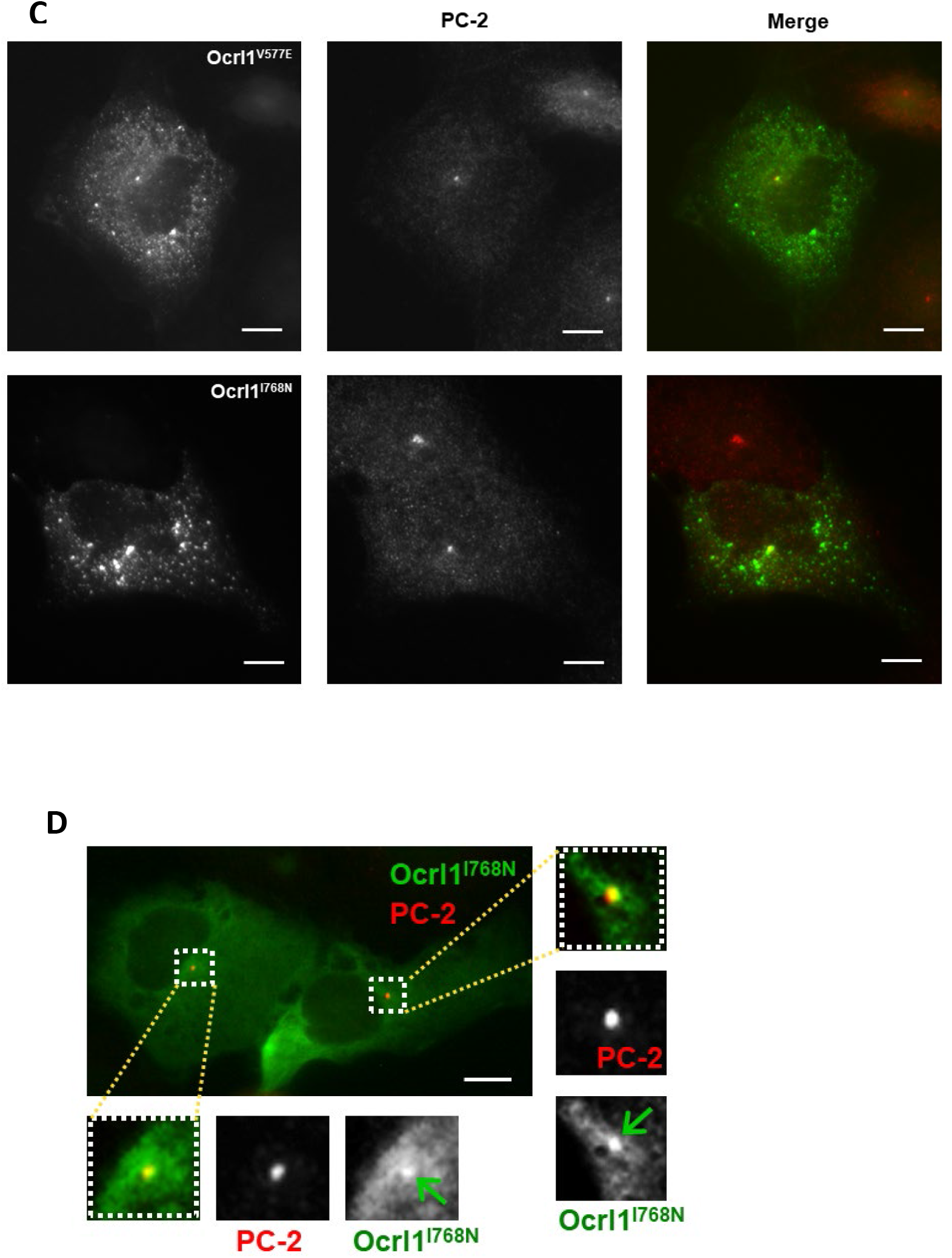
ASH-RhoGAP domain mutants exhibit enrichment or accumulation at the centriole under steady state conditions. **A**: HK2 KO cells transiently expressing Ocrl1^WT^ or ASH-RhoGAP mutants and maintained in complete media (left panel), were immunostained for the centriole marker PC-2 (see *Materials and Methods*). Arrows indicate Ocrl1 at PC-2 labeled structures. **B**: Transfected cells were randomly imaged from at least 25 fields containing at least 40 cells. Cells with Ocrl1 colocalization to PC-2 were scored and fraction of cells exhibiting colocalization in a field was determined. Statistically significance of the mean difference with respect to Ocrl1^WT^ was **p<(0.05/3=0.0167) (Bonferroni correction) by the Wilcoxon test; box plot of a representative experiment. Reference line indicates enrichment observed in GFP-transfected cells. **C**: HK2 KO cells transiently expressing Ocrl1^V577E^ or Ocrl1^I768N^ exhibiting protein aggregation. **D**: HK2 KO cells stably expressing Ocrl1^I768N^ immunostained for PC-2 (centriole marker) (see *Materials and Methods*). Highlighted region in merged images corresponding to the perinuclear region which was scaled to 3X (inset images) to better visualize Ocrl1 and PC2 colocalization. Arrows indicate Ocrl1 at PC-2 labeled structures. Scale bar: 10μm.

While HK2 KO cells expressing Ocrl1^WT^ showed a continuous and compact Golgi complex, as seen by labeling the *trans*-Golgi (using an anti-TGN46 antibody) and the *cis*-Golgi (using an anti-GM130 antibody, Supplemental Fig. 2) networks, cells expressing patient variants H524R, D451G and V508D displayed a discontinuous and fragmented Golgi complex that occupied a larger cellular area than the one covered by the GA in cells expressing Ocrl1^WT^ (Fig.5A, B and Supplemental Fig. 2). Interestingly, we also observed that these Ocrl1 mutated variants displayed a dispersed punctate pattern themselves with poor colocalization between the mutated Ocrl1 variant and TGN46 (Fig. 5A, insets) and lacked phosphatase activity (Fig. 5C).

Importantly, the magnitude of the GA fragmentation was independent of the amount of GFP-tagged Ocrl1 mutated variant expressed (measured as total cell-associated green fluorescence intensity, Supplemental Fig. 3). Along the same lines, this phenotype was also observed in HK2 KO cells stably expressing the 5’-phosphatase affected Ocrl1 variants H524R and D451G (Fig. 5D,E). Similar results were observed using HeLa and HEK293T cells (Data not shown).

In contrast, cells expressing Ocrl1^S256N^ did not show substantial GA fragmentation (Fig. 5A,B). Importantly, *in vitro* malachite green 5’ phosphatase assays using GST-Ocrl1^S256N^ (see *Materials and methods*) revealed that this variant retained 5’ phosphatase activity (Fig. 5C).

#### c) ASH-RhoGAP mutated patient variants show mislocalization to centriolar structures

Expression of GFP-tagged ASH-RhoGAP mutants Ocrl1^V577E^, Ocrl1^F668V^ and Ocrl1^I768N^ in HEK293T KO cells under steady state conditions displayed a characteristic perinuclear punctate structure, same observations were made using HK2 KO cells (Fig.6A).

A similar localization was previously observed for Ocrl1^WT^, but *ONLY under ciliogenesis induction conditions (i.e., serum-starvation*) (15). Specifically, we have shown that during ciliogenesis, Ocrl1^WT^ localize transiently at the base of the cilium (where the axoneme-linked centrioles localize), presumably for trafficking of ciliary cargo to the PC (15). Therefore, and *although steady state experiments (like the ones shown in Fig. 6A) are not done under serum-starvation conditions*, we wondered if the punctate structure observed in cells expressing ASH-RhoGAP mutated Ocrl1 would also correspond to the centrioles. Indeed, we found that GFP-tagged Ocrl1^V577E^, Ocrl1^F668V^ and Ocrl1^I768N^ perinuclear punctate structures colocalized with pericentrin-2 (PC2: a centriole marker, Fig. 6A,B). In contrast, under the same conditions (*using serum-supplemented media; e.g*., Fig. 6A), we could *not* observe substantial localization of Ocrl1^WT^ at the centriole.

In addition, ASH-RhoGAP mutated Ocrl1^V577E^, Ocrl1^F668V^ and Ocrl1^I768N^ patient variants had an overall diffuse cytosolic appearance in cells and lacked typical perinuclear enrichment at the Golgi apparatus. We also analyzed centriolar localization of the ASH-RhoGAP mutated proteins as function of their levels of expression. Specifically, we segregated transfected cells into ‘low’ (>80% of the total cell population), ‘medium-high’ total Ocrl1 variant content (*i.e*., total fluorescence intensity) groups covering the entire cell population (Supplemental Fig. 4A, B). Independently of the total fluorescence intensity, GFP-tagged Ocrl1 patient variants Ocrl1^V577E^, Ocrl1^F668V^ and Ocrl1^I768N^ colocalized with the centriole (Supplemental Fig. 4A). Further, at higher levels of expression, Ocrl1 mutated proteins Ocrl1^V577E^ and Ocrl1^I768N^ showed evidence of protein aggregation (Fig. 6C and Supplemental Fig. 4B). Although this may not occur at physiological levels of expression, it suggested a destabilizing effect of these ASH-RhoGAP mutations on Ocrl1.

Importantly, and in contrast to Ocrl1^WT^, low copy number, stably-transfected Ocrl1^I768N^ cells also showed centriolar and diffuse cytosolic localization lacking perinuclear enrichment (Fig.6D). In agreement with our observations on low-expressing transient transfectants, stably expressing cells did not exhibit protein aggregates (Fig. 6D, Supp. Fig.4B).

### Selected *OCRL1* mutations affecting phosphatase domain likely cause a conformational change in the catalytic domain of the mutated protein

Although some of the tested Ocrl1 patient variants exhibited changes in phosphatase domain’s residues not directly involved in binding/processing of substrate (Supplemental Fig. 1), they produced cellular defects linked to lack of enzymatic activity.

The phosphatase domain contains 6 conserved motifs essential for 5’ phosphatase activity (37–40), as well as other conserved residues involved in substrate binding and lipid chain interactions (39, 41) (Supplemental Fig. 1). For example, the mutation H524R affects a critical residue that interacts with the scissile 5’ phosphate (39, 41) (Supplemental Fig. 1), and has been demonstrated to abolish phosphatase activity (28). In contrast, D451G and V508D affect residues that are found outside the conserved catalytic motifs within the phosphatase domain (Supplemental Fig. 1), and yet were affecting cellular functions of Ocrl1 (Figs. 2,3,5; Supplemental Figs 2, 3). We hypothesized that *these mutations affect the conformation of the catalytic domain, impairing its phosphatase activity*.

We used molecular dynamics (MD), to model residue changes D451G and V508D in the phosphatase domain of Ocrl1 (PDB ID: 4CMN). Our results predicted that although the changed amino acids were physically distant from the enzyme’s active site, they induced conformation changes affecting critical catalytic residues (Fig.7A, B). Further, we calculated per residue root mean square fluctuation (RMSF) and root mean square deviation (RMSD), to quantify the conformational change induced by the site-specific mutations. RMSF (to identify flexible domains in Ocrl1) indicated that the D451G and V508D point mutations made the catalytic site comprising of residues in range 420 – 500 and 250 – 305 less flexible, respectively (Fig 7C, D). Analyzing RMSD, MD results indicated, that residues 250 – 305 in the catalytic site of Ocrl1 deviated with respect to WT for patient variants V508D and D451G (Fig. 7E, F).

**Fig. 7.**
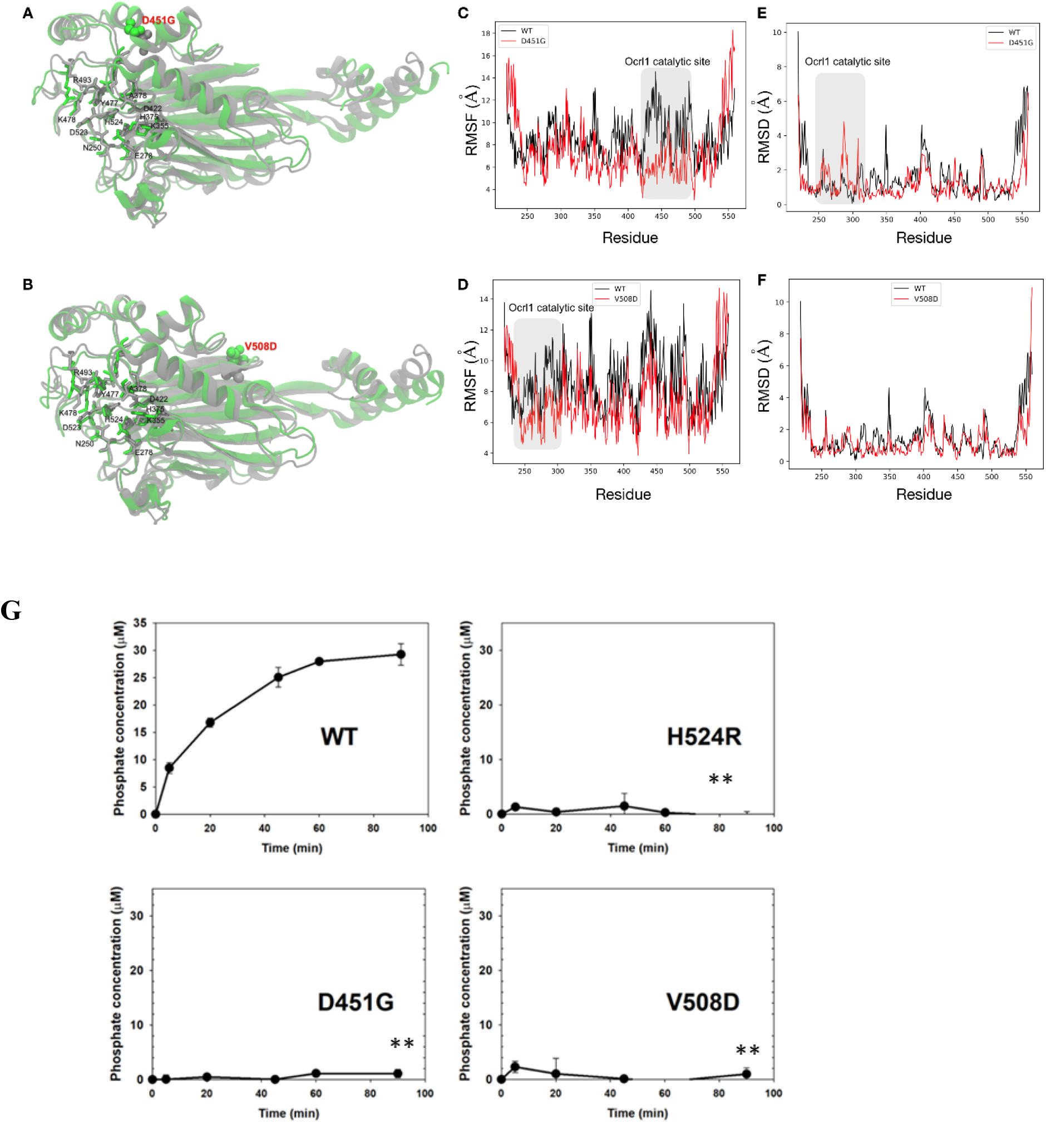
Molecular dynamics prediction of the effect of D451G/V508D mutations on Ocrl1 phosphatase domain structure. **A**: Conformational change in WT (gray) and mutant (green). The residues in catalytic site are represented by sticks, while the point mutation (D451G) is shown by a space-filled model. **B**: Conformational change in WT (gray) and mutant V508D (green). **C:** Root mean square fluctuation (RMSF), comparison between WT and mutant D451G, suggesting that the active site residues in range 420-500 becomes less flexible. **D:** Root mean square fluctuation (RMSF), comparison between WT and mutant V508D, suggesting that the active site residues in range 250-305 becomes less flexible. **E:** Root mean square deviation (RMSD) per residue for WT and mutant D451G, indicating that the catalytic domain (residues 250 −305) deviates around 2 Å. **F:** Root mean square deviation (RMSD) per residue for WT and mutant V508D, indicating that the residues in the catalytic domain of Ocrl1 do not deviate significantly with respect to wild-type. **G**: Ocrl1 phosphatase domain mutants are impaired for 5’ phosphatase activity. Bacterially expressed and purified GST-Ocrl1^1-563^; WT (top panel left) and the different phosphatase mutants were assayed for enzymatic activity *in vitro* using the malachite green method. Experiments were done at constant enzyme and substrate concentrations while varying the incubation times. All experiments were done in triplicates and repeated at least thrice. Statistically significance of the mean difference with respect to Ocrl1^WT^ at the latest time point was **p<(0.05/3=0.0167) (Bonferroni correction) by Student t-test.

Although results shown in Fig. 5C already showed that mutations D451G and V508D affected Ocrl1 phosphatase activity, we performed a more detailed analysis of the catalytic function of these variants using *in vitro* malachite green phosphatase assays (*Materials and Methods*) and bacterially purified GST-fusions of Ocrl1^1-563^ (*i.e*., truncations containing the PH-phosphatase domain) from patient variants (S256N, D451G, V508D and H524R) or WT.

As expected, Ocrl1^H524R^ lacked phosphatase activity (Fig.7G). Importantly, and in agreement with our MD results, although the D451G and V508D amino-acid changes do not directly affect the catalytic site of Ocrl1, they led to 5’-phosphatase domain inactivation (Fig.7G).

## DISCUSSION

Complementing the pioneering works of different authors (12–16, 28, 31, 42–49) this study contributes to better understand LS as a complex disease. Here we highlight the heterogenous nature of this condition by focusing on the impact of different *OCRL1* mutations on typical cellular phenotypes while reporting previously unnoticed LS abnormalities triggered by specific patient variants. Indeed, to the best of our knowledge, this is the first systematic analysis of the specific phenotypic impact of a group of mutations encoding for residue changes in *all* regions of Ocrl1.

On the one hand, data presented in this study supports the hypothesis that some Ocrl1’s functional roles are segregated within the protein primary structure (12, 15, 46). Specifically, that integrity of the N-terminus is mostly required for membrane remodeling functions (*e.g*., cell spreading (12)), while Ocrl1’s C-terminal region is more relevant to ciliogenesis (15). As indicated before, the central phosphatase domain was required for both functions (12, 15).

On the other hand, this study highlights specific phenotypes associated with certain mutations.

A predicted Ocrl1 patient variant that lacks its N-terminal PH domain (Ocrl1^ΔPH^), but bears all other domains/regions should be able to interact with Rab GTPases and in consequence should have proper intracellular localization (50–52). Indeed, similar to Ocrl1^WT^ this variant was enriched in compartments like the TGN; however, we also observed Ocrl1^ΔPH^ cytosolic/nuclear mis-localization. These observations led us to hypothesize that absence of the PH domain led to the loss of previously unnoticed localization determinants, that allowed the GFP-tag to aberrantly drag/mislocalize the fusion protein to the nucleus. Nuclear localization is typical of free GFP (see Fig. 4 top panel), but it is not observed in GFP-tagged Ocrl1^WT^ where the protein’s strong localization determinants overpower the tag tendency to accumulate in the nucleus. Since Ocrl1^ΔPH^ lacks the N-terminal region that contains binding sites for the endocytic machinery, one could speculate of a possible role for proteins involved in endocytosis in sustaining Ocrl1 proper localization. Alternatively, this may be revealing the presence of yet to be identified Ocrl1-interaction partners in the N-terminus that play a critical role in maintaining its cellular localization. Further investigations will attempt to shed light into the mechanism by which the PH domain contributes to Ocrl1 proper localization. This variant produced cell spreading abnormalities in agreement with the previously proposed segregation of Ocrl1’s functions across the protein domains (12, 15).

Localization was also affected for Ocrl1 patient variants bearing amino acid changes within the ASH-RhoGAP domains. Specifically, we found that these mutants were predominantly cytosolic diffuse and lacking the Ocrl1^WT^-like enrichment at the GA. For these variants we also observed aggregates in a dose-dependent manner. Indeed, mutations affecting this domain are predicted to be destabilizing in nature, resulting in aggregated and/or degraded Ocrl1 (43, 53).

The presence of the ASH domain in other proteins is associated with localization at the centriole and cilia (11). Indeed, it is within this region that Ocrl1contains Rab-binding sites (50, 51) that during starvation (ciliogenesis-stimulating conditions) contribute to localization at the base of and support the assembly of the primary cilia (15). However, Ocrl1 patient variants with their ASH-RhoGAP domains mutated localized at centriole *even under steady state conditions* (which was not the case for Ocrl1^WT^). Importantly, this was observed even in a Rab-binding mutant Ocrl1^F668V^ suggesting that this phenomenon is likely related to unique characteristics that these mutants possess.

Although these mutants produced aggregates, it is unclear if this phenomena occur in patients as it was most noticeable at higher levels of expression. Interestingly, in these cases we observed aggregates colocalizing with centrioles (Fig. 6D) and it is well-known that subunits and substrates of the proteosome are enriched at these structures (54). Given the proposed unstable nature of ASH-RhoGAP mutants, it is tempting to speculate that this enrichment is due to unstable proteins being targeted for degradation by the centriole-associated proteasome known as the ‘aggresome’ (55–57). In fact, many other aggregated proteins implicated in diseases have been observed to be enriched in these compartments as well (55, 57).

Several patient variants with mutated phosphatase domains (affecting residues with either catalytic and non-catalytic relevance) displayed a dispersed punctate pattern and induced Golgi apparatus fragmentation. Further, these Ocrl1 puncta colocalized poorly with the GA. It should be noted that Golgi complex fragmentation has been observed in neurological diseases (24–26), and therefore might be relevant for the neurological component of LS.

GA fragmentation is only dependent on lack of Ocrl1 catalytic activity; therefore, it is possible that PI(4,5)P2 accumulation affects GA integrity. Interestingly, the Nussbaum lab previously reported that Ocrl1 interacts with the GA tethering protein Golgin-84 (58), and other studies have demonstrated that trafficking defects of the latter led to GA fragmentation (59–61). It is possible that, in the absence of Ocrl1 catalytic activity, Golgin-84 is not properly localized and causes the GA phenotype.

In addition, these mutants produced substantial defects in cell spreading and ciliogenesis further supporting the idea that loss of enzymatic function affects both cellular processes. Interestingly, the LS patient variant Ocrl1^S256N^, despite of having a mutation in the phosphatase domain, had a substantial phosphatase activity and in consequence did not induce GA fragmentation or displayed a dispersed punctate pattern (Fig. 5A, B, D). However, while cell spreading of cells expressing this variant was normal, Ocrl1^S256N^ could not sustain normal ciliogenesis (Figs. 2B and 3A, respectively). Therefore, the Ocrl1^S256N^ variant may cause LS by an almost exclusively ciliogenesis-dependent mechanism and is the focus of further investigations.

We humbly believe that one of the most interesting hypotheses emerging from this work is that LS has a conformational disease component; *i.e*., that *some* patients would express Ocrl1 variants which are conformationally affected.

In fact, out of the 80 unique missense mutations within the phosphatase domain of Ocrl1, ~50% of the mutations (including D451G and V508D) are affecting residues not directly involved in catalysis and yet produce LS. In other words, we speculate that some patients express Ocrl1 mutated proteins with intact binding/catalytic sequences, but likely locked in a conformation unable to process substrate. Indeed, the lack of phosphatase activity of some of these mutants and molecular dynamics analysis supported our rationale (Fig. 7).

Therefore, we speculate that for patients expressing these specific variants, LS has a component of a ‘conformational or misfolded protein disease’. A conformational disease is one where mutations in the gene lead to loss of function in the resultant mutant protein, either by affecting synthesis, transport stability, protein folding or its enzymatic activity (62). These abnormalities cause the accumulation of the non-native conformation, rendering loss in function (62, 63). This possibility is the focus of current and intense investigation.

Overall, this study, along with others, suggests the existence of several levels of complexity/variability contributing to make LS a disease with a broad spectrum of symptoms and phenotype severity among patients.

At tissue-cellular level, the severity or manifestations of Ocrl1 functional deficiency would depend on the levels of expression of the Ocrl1’s paralog Inpp5b in different cell types/tissues/organs (48). However, as is to be expected for different gene products, even in the presence of Inpp5b, Ocrl1-deficiency triggers Ocrl1-specific phenotypes (*e.g*., cell spreading and migration; (12)). Nevertheless, we speculate that different cell types/tissues/organs would show various levels of severity of these specific phenotypes depending on factors like composition and compliance of the extracellular matrix. As a result of this composite of factors different organs would be affected in different manner by *OCRL1* mutations.

The next level of variability affecting patients comes from the *nature* of the specific LS-causing *OCRL1* mutation: missense, nonsense or deletion/insertion mutations (with the latter in either coding or non-coding regions of the gene) and their possible outcomes (*e.g*., presence of a mutated protein or absence of gene product, splicing defects and changes in expression levels of Ocrl1). This study focused on different missense mutations affecting different Ocrl1 domains or producing a ΔPH-truncated variant (31) and their effect on cellular phenotypes.

Our results support the idea of spatial segregation of functions for the Ocrl1 molecule: the N-terminus required for membrane remodeling, the C-terminus for ciliogenesis and the central phosphatase domain for both processes. Therefore, *OCRL1* mutations affecting these different regions have different impact on cellular phenotypes (Fig. 8); adding another source of variability to LS patient’s spectrum of phenotypes and symptoms. Further, for different LS-causing mutations, the severity of the corresponding phenotype was not homogeneous even when affecting the same domain (Fig. 8). In addition, mutations led to domain-specific abnormalities such as, mislocalization or Golgi apparatus fragmentation (Fig. 8).

**Fig. 8.**
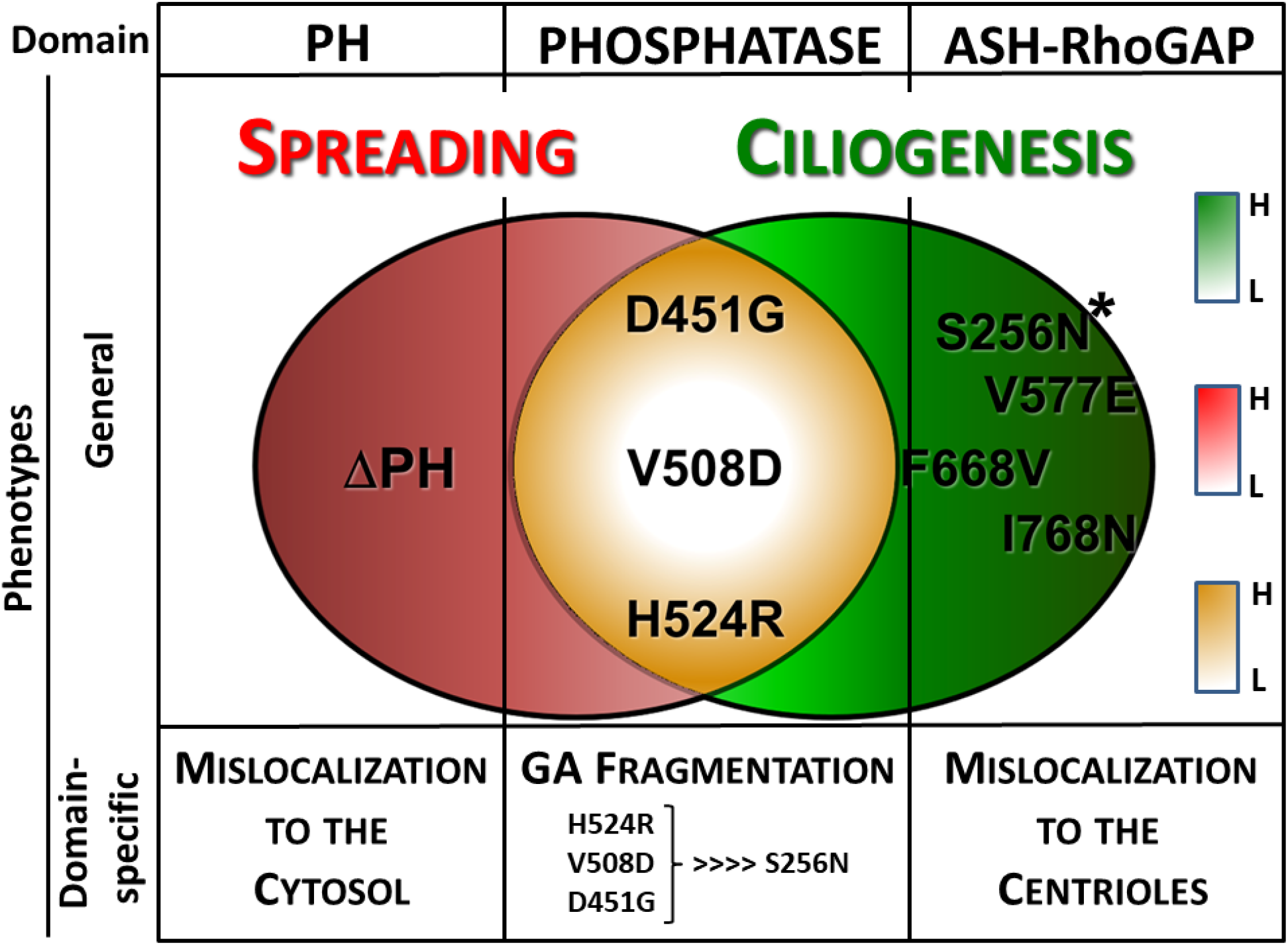
Different missense mutations effect on LS phenotypes. Specific mutations have differential effects on general LS phenotypes such as defects on cell spreading (*red*) and ciliogenesis (*green*). Different mutations associated with specific amino acid changes (or alternative protein initiation, *e.g*., ΔPH) at specific Ocrl1 domains are indicated. Venn diagrams show patient variants inducing only cell spreading defects (ΔPH), only ciliogenesis abnormalities (e.g., ASH-RhoGAP mutated variants) or both (phosphatase mutated proteins, in *yellow*) Differential phenotype severity is represented by color tone (boxes on the right), the darker the tone the more severe the phenotype, while *white* indicates the mildest phenotype. Phenotypes associated with changes in specific Ocrl1 domains are indicated. *: patient variant inducing severe (but only) ciliogenesis phenotypes.

This study provides evidence for heterogeneity in LS cellular phenotypes indicating that the impact of *OCRL1* mutations is a complex composite of all the above factors. In addition, genetic modifiers as well as environmental factors may play a role in how these defects manifest as patient symptoms and should be the focus of further studies.

## Materials and Methods

### Reagents and constructs

All reagents were procured from Fisher Scientific (Fairlawn, NJ) or Sigma Aldrich (St. Louis, MO) unless stated otherwise. Antibodies used in this study are listed in Supplemental Table I. Site-directed mutagenesis was performed using Quikchange Lightning mutagenesis kit (Agilent Technologies) and pEGFP-c1 hs*OCRL1* (wild-type, isoform b) as template to create the various *OCRL1* constructs used in this study. Plasmid constructs used in this study are listed in Supplemental Table II.

### Cells and culture conditions and transfections

Normal human proximal tubule epithelial (HK2) and human embryonic kidney epithelial 293T (HEK293T) cells were purchased from ATCC and cultured in DMEM, Streptomycin/Penicillin, 2mM L-Glutamine and 10% fetal bovine serum (FBS). Cells were maintained at 37 °C in a 5% CO_2_ incubator. *OCRL1^-/-^*(*OCRL* KO) HK2 and HEK293T cells were prepared by GenScript Inc. Piscataway, NJ, USA and maintained under identical conditions. Absence of Ocrl1 as well LS-specific phenotypes including cell spreading and ciliogenesis defects were previously validated in the KO cell lines (46, 64). Plasmids encoding different Ocrl1 mutants were transfected using Fugene 6 reagent (Promega) according to manufacturer’s instructions.

### Preparation of stable cell lines

HK2 (and HK2 KO) cells have been immortalized using a plasmid encoding SV40 large T antigen than confers it resistance to G418. Therefore, to prepare stable transfectants cells were co-transfected with plasmid for expression of GFP-Ocrl1^WT/Mutants^ along with pTre2Hyg-6XHis vector (in a 4:1 ratio). Stably transfected clones were selected using 100ug/ml Hygromycin (Bruce et al.). Antibiotic-containing media was replaced every 48h. Within 2-3 weeks, GFP positive colonies were identified and isolated using sterile cloning discs. These clones were sub-cultured over multiple passages and expression of Ocrl1^WT/Mutant^ was confirmed by microscopic analysis.

### Cell spreading assays

Cells were transfected as described above and grown in complete media upto 18h. Cell confluency was maintained at ~50% to ensure single cell suspensions were obtained prior to spreading assays. 18h post transfection, the cells were lifted with 20mM EDTA (in 1X PBS), pelleted at 100xg for 5min and resuspended in complete media. Cell suspensions were then set in a rotator for 1 hour before seeding them on 10ug/ml fibronectin-coated coverslips for 30min, undisturbed, to allow attachment and spreading. At 30min, coverslips were gently washed using 1X PBS and fixed in 4% formaldehyde for 10min at room temperature. Cells were stained with rhodamine–phalloidin (used at 1:200) and imaged by epifluorescence microscopy. At least 40 cells were analyzed per experiment. The magic wand tool in ImageJ software was used to trace the cell boundaries and determine cell areas and perimeter.

In every experiment, spreading area of cells expressing Ocrl1 mutants was normalized to the median area calculated in Ocrl1^WT^-expressing cells for the same experiment. Histograms were constructed from the normalized spreading area values and Kolmogorov-Smirnov (KS) test was performed to determine statistical significance.

### Ciliogenesis assays

Cells were seeded (in complete media) on glass cover slips coated with poly-D-lysine and transfected as described above. Confluency of cells was maintained in such a way that it did not exceed 50% at the time of ciliogenesis. 18h after transfection, the media was replaced by 0.1% serum DMEM (starvation media) for another 24h to induce ciliogenesis. Cells were washed with 1X PBS, fixed in 4% formaldehyde–PBS for 10min. Indirect immunofluorescence was performed using antibodies against acetylated tubulin antibody (to label cilium) and pericentrin-2 (PC-2; to label centriole) (refer to Table I). 20 random fields, comprising of at least 50 cells were imaged for every experiment and repeated at least thrice.

From all the transfected cells imaged, the fraction of Ocrl1 mutant-transfected cells forming a cilium was calculated. This was normalized to the fraction of cilia produced by Ocrl1^WT^-expressing cells from the same experiment. The upper limit of fraction of ciliated cells was 1 (represented by HEK293T KO cells expressing Ocrl1^WT^) while the lower limit of fraction of ciliated cell was 0.7 (HEK293T KO cells expressing Ocrl1^GFP^). Though we observed variability between experiments, individual experiments were statistically significant.

### Indirect immunofluorescence and fluorescence microscopy

In all immunofluorescence procedures, antibodies were diluted in DMEM media containing 10% FBS (blocking agent) and 0.1% saponin (permeabilizing agent). Primary antibodies were incubated for 1h at room temperature, washed 2X with PBS. Fluorescent molecule-conjugated secondary antibodies were incubated with cells for 45min in the dark, washed 2X with PBS. Cells were then stained using DAPI to label the nucleus and mounted on pre-cleaned glass slides using Aqua-PolyMount reagent (Polysciences). Following indirect immunofluorescence, coverslips were imaged with constant fluorescence exposure times using a 40X objective on Zeiss Axiovert inverted microscope. Random fields were imaged to cover the entire coverslip. Exposure times were maintained consistent for all independent experiments.

### Protein purification

The PH and phosphatase domain of wild-type hsOcrl1 (1-563 amino acids) was cloned in a pGEX-4T1 plasmid that contains an N-terminus glutathione S-transferase (GST) tag. Using site directed mutagenesis, missense mutations H524R, D451G and V508D were introduced. Plasmids were transformed in Rosetta (DE3) competent cells. Bacterial cultures were grown overnight at 37°C, in LB medium supplemented with 2.5% glucose, 1X ampicillin, and 1X chloramphenicol. The following day, cultures were expanded in super broth media containing 1X ampicillin and 1X chloramphenicol for 3 hours at 37 °C. Then, cultures were supplemented with 5% glycerol and 0.1 mM IPTG and incubated for 5 hours at 30 °C.

Cells were harvested by centrifugation (3,000 x g, 10 min), and pellets were stored at −80 °C until use. Cells were lysed in lysis buffer containing 200 mM Tris pH 7.4, 10% glycerol, 01.% Tween 20, complete EDTA-free protease inhibitor, 1 mg/ml lysozyme was added to resuspend prepared cell pellets. Cells were disrupted by sonication at 50% power for 3 sets of 33 pulses with 30 second breaks in between pulses. Cell debris was removed via centrifugation at 21,500 x g, 30 min, 4 °C. The supernatant was transferred to tubes containing glutathione resin (Pierce) and incubated at room temperature on a shaker (12 rpm) for 2 hours. Beads were washed 4 times with lysis buffer (w/o protease inhibitor and lysozyme). Then, 100 mM glutathione (pH 8.0) was added to the tubes containing supernatant and incubated at room temperature on a shaker (12 rpm) for 2 hours to elute the protein. Supernatant (after centrifugation at 1,000 x g for 2 min) was loaded onto desalting columns (Thermo Scientific, Zeba, 89891) and centrifuged (1,000 x g, 2 min, acceleration: 5) and purified protein was obtained. Protein concentration was estimated using NanoDrop 1000 (ThermoFisher) and was used immediately for malachite green phosphatase assays.

### Malachite green phosphatase assays

For 5’ phosphatase activity assays, the malachite green phosphate assay kit (Sigma-Aldrich, MAK307) was used. Briefly, in a 384-well plate, 10 μl of 2μM of purified Ocrl1 (wildtype and mutant) was incubated with drug or vehicle for one hour at room temperature, to obtain a final enzyme concentration of 1uM. After treatment,10 μl of 50 μM PI(4,5)P2 diC8 (Echelon Biosciences, P-4508) was added to wells and incubated for 5 minutes at room temperature in the same buffer as the purified phosphatases, containing 200 mM Tris, pH 7.4. To stop enzyme reaction, 20 μl of 0.25X malachite green reagent was added to the reaction wells. After 20 minutes of color development, absorbance was measured at 620 nm. A standard phosphate curve was prepared (as per manufacturer’s instructions) in the enzyme buffer solution to determine the amount of free phosphate released by the enzyme variants tested. Experiments were repeated at least thrice and each condition was tested in triplicates. Student’s t-test was used to determine statistical significance.

### Molecular Dynamics (MD) simulation

For modeling a bound complex for Ocrl-1 (PDB: 4CMN) (39) and PIP2, the protein is constructed with explicit lipid bilayer with membrane composition 90% phosphatidylcholine (PC), 5% phosphatidylserine (PS) and 5% PIP2 (65) using CHARMM-GUI membrane builder (66, 67) (Supplemental Fig. 5).

The protein and membrane systems are then solvated with TIP3P water and neutralized with 200 mM MgCl_2_. MD simulations are performed for 100 ns to obtain an equilibrated model for WT Ocrl-1 (enzyme) and PIP2 (substrate) complex. Supplemental Fig. 5B shows the trajectory for the head group atoms of PIP2 molecule that interacts with the residues in catalytic site of Ocrl1 (WT), starting from the membrane plane (red), diffusing inwards into the cytoplasmic region (blue).

Next, the enzyme-substrate complex is used, and point mutations for D451G and V508D are performed using the MUTATOR plugin (https://www.ks.uiuc.edu/Research/vmd/plugins/mutator/) in VMD 1.9.3 (68). Independent MD simulations, each 100 ns long (in addition to equilibration) are performed for WT and mutant (D451G and V508D) systems.

All the MD simulations were performed with NAMD 2.13 (69) using CHARMM36m force field for lipid/protein (70) and a timestep of 2 fs. Long range electrostatic interactions were evaluated with particle mesh Ewald (PME) (71) and periodic boundary conditions were used throughout the simulations. Non-bonded forces were calculated with a cutoff of 12 Å and switching distance of 10 Å. During the simulation, temperature (T = 310 K) and pressure (P = 1 atm) (NPT ensemble) was maintained by Nosé-Hoover Langevin piston method (72).

### Rosetta (ΔΔG) calculations

This calculation predicts the change in stability (ΔΔG) of the enzyme induced by a point mutation using in ddG_monomer application in ROSETTA (73). The application takes as input the crystal structure of WT (pre-minimized) and generates a structural model of the point-mutant. The ΔΔG is given by the difference in Rosetta energy between the WT and the point mutant structures. Specifically, 50 models each of the WT and mutant structures are generated, and the most accurate ΔΔG is taken as the difference between the mean of the top-3-scoring WT and mutant structures. Negative ΔΔG values indicate increased stability, where ΔΔG = mutant energy - wildtype energy (Supplemental Table III).

### Statistical analysis

Statistical significance of differences between spreading-distribution histograms were analyzed using the Kolmogorov-Smirnov (KS) test. The student’s t-test was used to evaluate the significance of differences of normally distributed samples (*e.g*., ciliogenesis experiments), while the Wilcoxon’s test was employed when samples were non-normally distributed (Quantification of Golgi apparatus fragmentation and PC2 colocalization). For all comparisons involving Ocrl1^WT^ and Ocrl1 mutants, the Bonferroni’s correction for multiple comparisons was performed whenever applicable [αC=p/n; n being the number of comparisons].

After carefully analyzing each data set distribution the most appropriate representation of data in each case was adopted. These representations included histograms and box plots as they allow to thoroughly examine the data distribution (74). When the data presented a normal distribution, a bar graph with standard deviations was used to represent the data.

## ACKNOWLEDGEMENTS

We are indebted to Drs. Donna Fekete, Don Ready and Phil Low (Purdue University) for stimulating discussions. We also thank members of the Aguilar lab for discussions and critical reading of the manuscript.

## Supplemental Material

**Supplemental Table I:** Antibodies used in this study

| Supplemental Table I: Antibodies used in this study |  |  |  |
| --- | --- | --- | --- |
| Antigen | Host | Source | Dilution used for Indirect Immuno-Fluorescence |
| Acetylated-Tubulin | Mouse | Sigma Aldrich (6-11B-1) | 1:1000 |
| Pericentrin-2 (PC-2) | Rabbit | Abcam (ab4448) | 1:300 |
| TGN46 | Rabbit | Thermo Fisher (PA5-23068) | 1:400 |

**Supplemental Table II:** Plasmids used in this study

| Supplemental Table II: Plasmids used in this study |  |  |
| --- | --- | --- |
| Plasmid name | Description | Source |
| GFP | pEGFP-c1 empty vector (expresses GFP) | Clontech |
| Ocr11<sup>WT a</sup> | GFP-Ocr11<sup>1-893</sup> in pEGFP-c1 | Coon et al. 2009 |
| Ocr11<sup>S256N</sup> | GFP-Ocr11<sup>1-893, S256N</sup> in pEGFP-c1 | This study |
| Ocr11<sup>D451G</sup> | GFP-Ocr11<sup>1-893, D451G</sup> in pEGFP-c1 | This study |
| Ocr11<sup>V508D</sup> | GFP-Ocr11<sup>1-893, V508D</sup> in pEGFP-c1 | This study |
| Ocr11<sup>H524R</sup> | GFP-Ocr11<sup>1-893, H524R</sup> in pEGFP-c1 | Coon et al. 2009 (12) |
| Ocr11<sup>V577E</sup> | GFP-Ocr11<sup>1-893, V577E</sup> in pEGFP-c1 | This study |
| Ocr11<sup>F668V</sup> | GFP-Ocr11<sup>1-893, F668V</sup> in pEGFP-c1 | This study |
| Ocr11<sup>I768N</sup> | GFP-Ocr11<sup>1-893, I768N</sup> in pEGFP-c1 | This study |
| Ocr11<sup>ΔPH</sup> | GFP-Ocr11<sup>187-893</sup> in pEGFP-c1 | This study |
| pTre2HygZ<sup>b</sup> | Used to confer resistance to hygromycin | Clontech |
| **a:** Human Ocr11 cDNA corresponding to isoform b (Coon et al., 2009) was used for all Ocr11-based constructs. **b:** used for co-transfection and selection of stable-transfected clones. |  |  |

**Supplemental Table III:** Stability (ΔΔG) for Ocrl1 point mutant with respect to WT

| Supplemental Table III: Stability ($\Delta\Delta G$)<br>for Ocr11 point mutant with respect to WT | |
| --- | --- |
| Mutation | $\Delta\Delta G$ (REU) |
| D451G | 7.298 |
| V508D | 10.899 |

### Supplemental Figures

**Supplemental Fig. 1.**
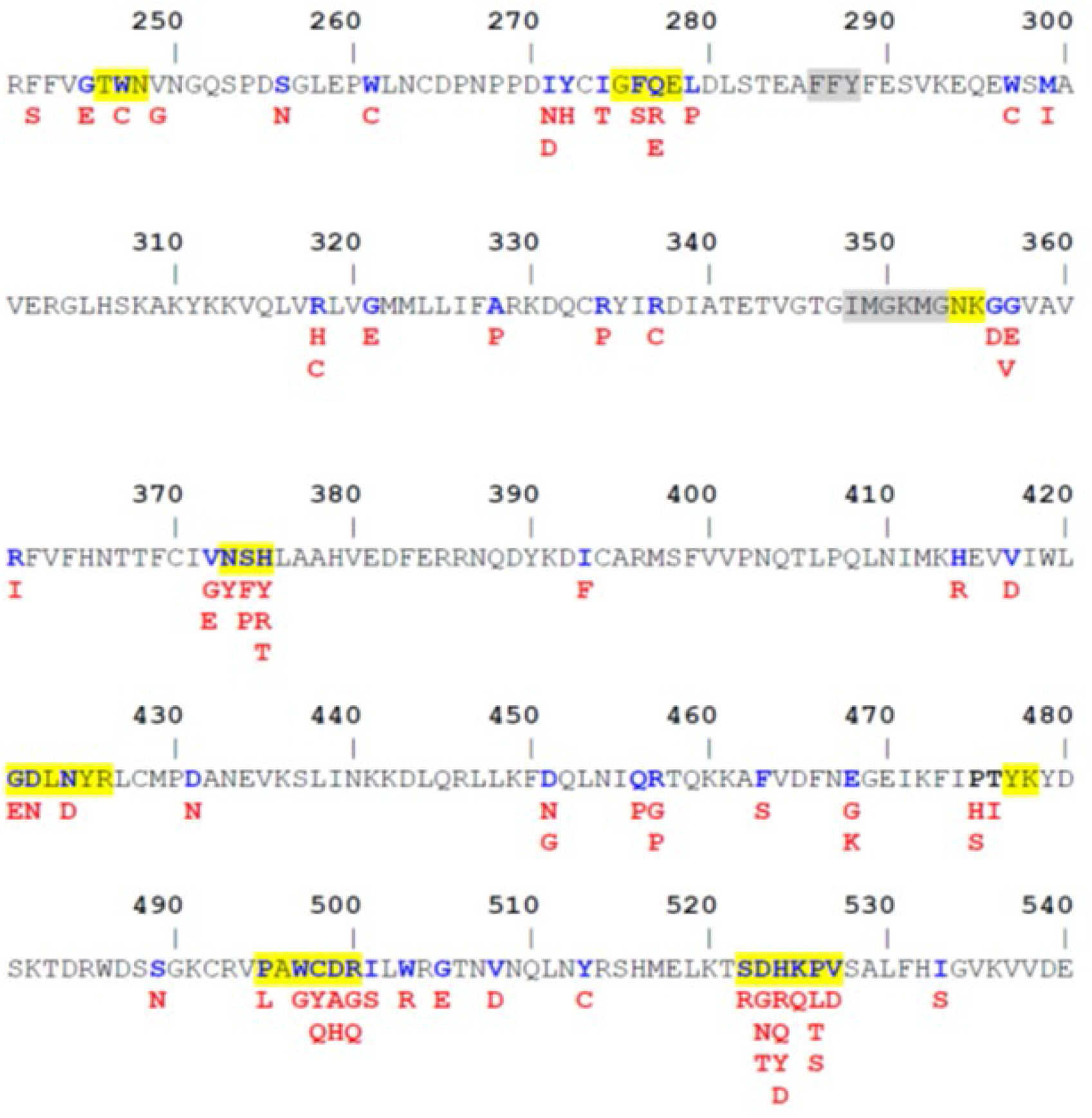
LS-causing mutations within the 5’-phosphatase domain. The amino acid sequence of Ocrl1 5’-phosphatase domain shows in blue residues affected by missense mutations in LS patients and in red the amino acids resulting of such mutations. Yellow rectangles enclose regions involved in binding/enzymatic processing of the phosphatidyl inositol moiety, while those in grey highlight carbon chain binding regions.

**Supplemental Fig. 2.**
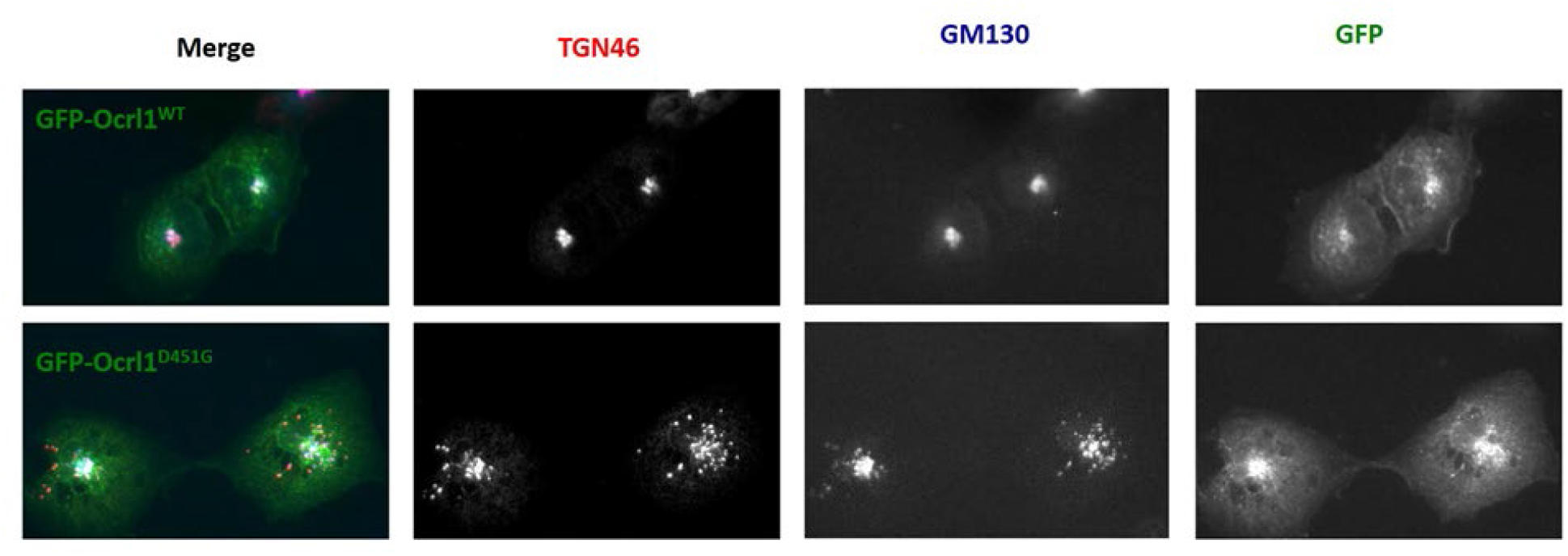
Ocrl1 patient mutations affecting the phosphatase domain cause Golgi Apparatus (GA) fragmentation. HK2 cells were transfected with Ocrl1^WT^(top) or Ocrl1^D451G^ (bottom), fixed and immunostained for TGN46 and GM130, markers for TGN (*trans*-Golgi network) and CGN (*cis*-Golgi network), respectively. Compared to WT-expressing cells, patient variant D451G-expressing cells exhibited a fragmented TGN and CGN that colocalize. Scale bar: 10μm.

**Supplemental Fig. 3.**
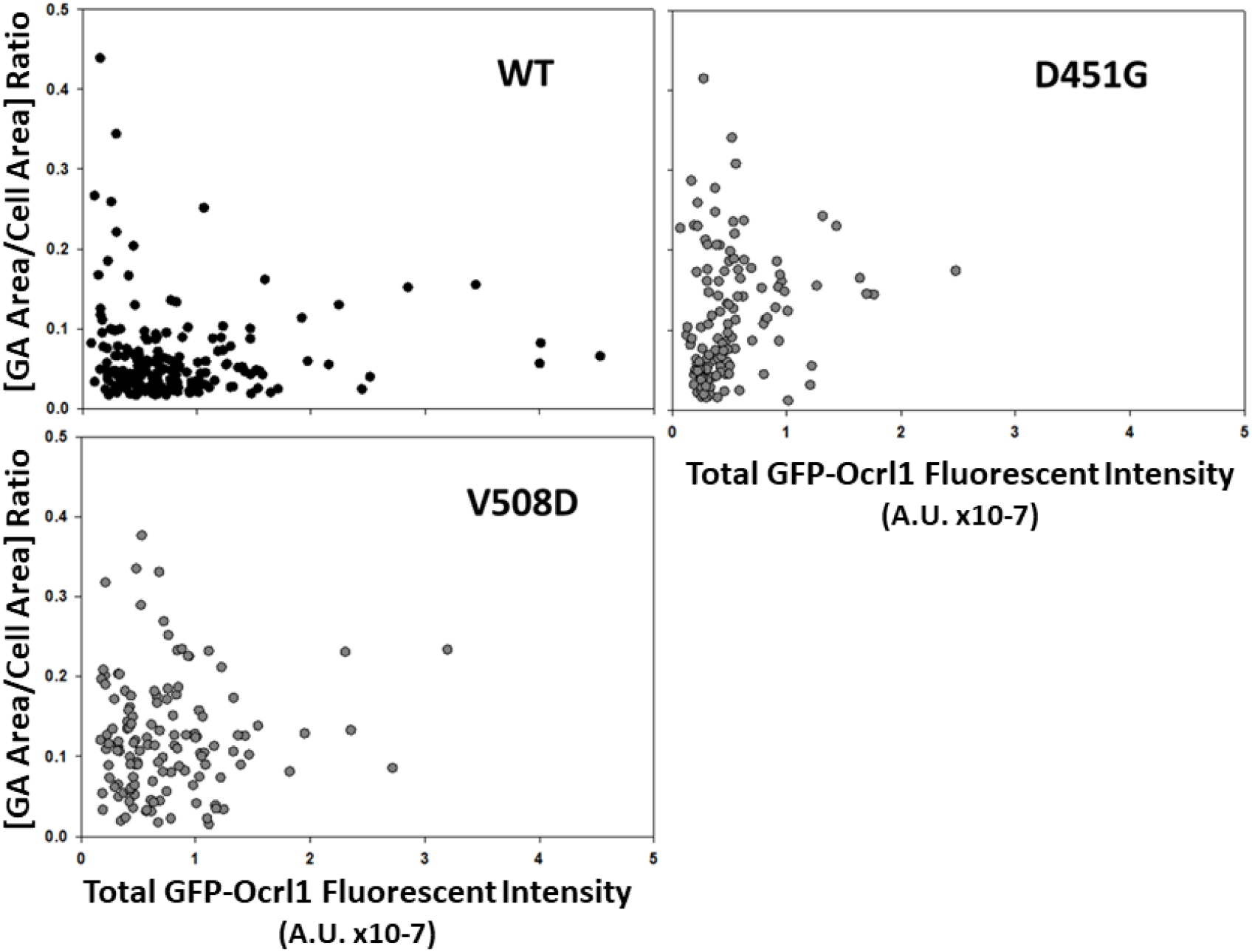
GA fragmentation is independent of the amount of Ocrl1 patient variant expressed. HK2 KO cells were transfected with plasmids encoding GFP-tagged Ocrl1^WT^ or phosphatase domain mutants. Following fixation and immunofluorescence using antibodies against TGN46, TGN area was measured and quantified using ImageJ. Simultaneously, total GFP fluorescence of transfected cells was measured using ImageJ and plotted against the corresponding TGN area/total cellular area. N=3 independent experiments, 120 cells/group.

**Supplemental Fig. 4.**
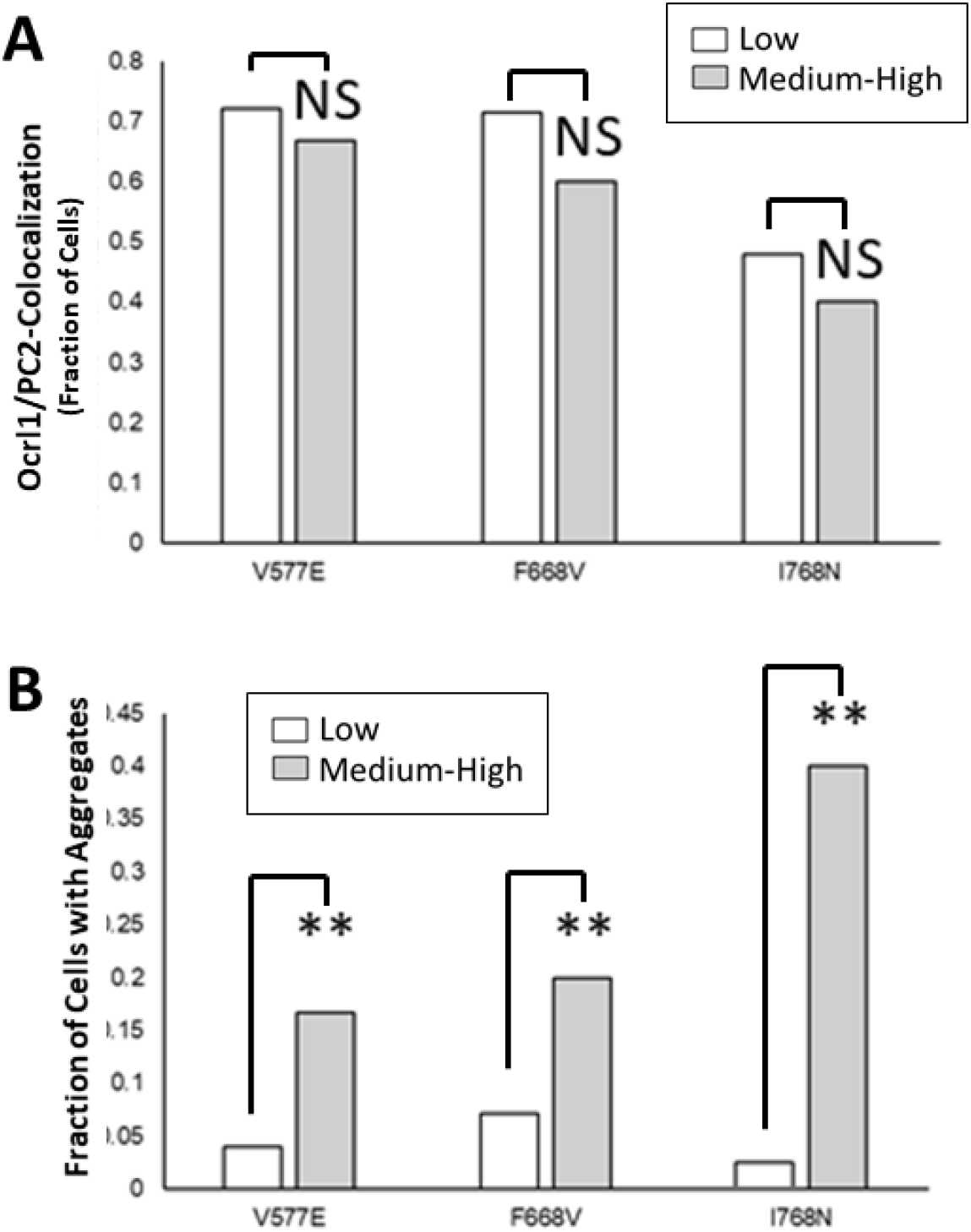
Ocrl1 ASH-RhoGAP mutants colocalize with centriole marker PC2 independently of patient variant levels of expression and aggregate in a dose-dependent manner. HK2 KO cells were transiently transfected with plasmids encoding GFP-tagged Ocrl1^WT^ or ASH-RhoGAP domain mutants. Transfected cells were then separated into 2 groups ‘low’ and ‘medium-high’ based on total GFP fluorescence intensity. **A.** Following fixation and immunofluorescence using antibodies against PC2 (pericentrin-2) and fraction of transfected cells with showing GFP-Ocrl1 patient variant colocalizing with PC2 was determined. **B**. Fraction of transfected cells exhibiting GFP-Ocrl1 patient variant aggregates was quantified. Graphs show results from 40 cells/group, in representative experiments. Signifance of the difference between Low and Medium-high Ocrl1-expression levels, **: p<0.05; NS: non-significant by the t-test.

**Supplemental Fig. 5.**
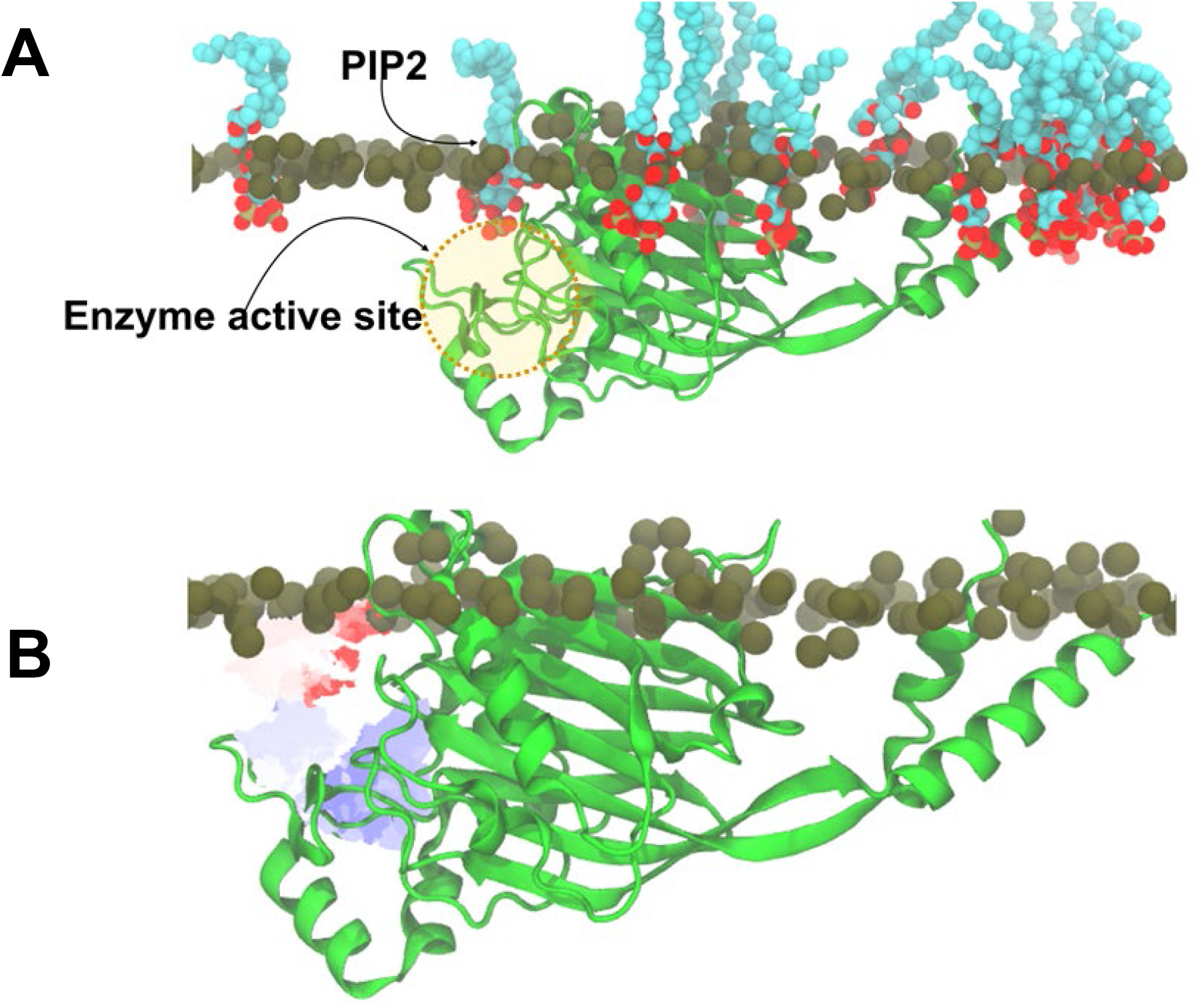
**A:** Molecular dynamics (MD) system of membrane bound *OCRL1* (PDB ID: 4CMN) represented in cartoon, substrate phosphatidylinositol 4,5-bisphosphate (PIP_2_) represented in space-filling. Only the phosphate atoms of other lipid molecules phosphatidylcholine (PC) and phosphatidylserine (PS) are shown as spheres, indicating the membrane plane. The active site in *OCRL1* is highlighted in yellow with a PIP_2_ molecule near the catalytic domain. **B:** Trajectory path of a single PIP_2_ molecule (labelled in panel B) from MD simulation. Red to blue shows the conformational transition from initial to final positions of the PIP_2_ substrate near and in the catalytic site during MD. The initial position of the protein and membrane planes are shown to provide a frame of reference for the PIP_2_ trajectory path.

